# Single-cell, spatial, and fate-mapping analyses uncover niche dependent diversity of cochlear myeloid cells

**DOI:** 10.1101/2024.10.30.621184

**Authors:** Aude Chiot, Max J. Felgner, Dillon Brownell, Katherine H. Rott, Alina Bogachuk, Dennis-Dominik Rosmus, Takahiro Masuda, Audrey Ching, Patrick J. Atkinson, Marco Prinz, Karen Sachs, Alan G. Cheng, Peter Wieghofer, Bahareh Ajami

## Abstract

Recent advances in fate mapping and single-cell technologies have revealed how the dynamics and function of tissue-resident macrophages are shaped by their environment. However, macrophages in sensory organs such as the cochlea where the central nervous system and peripheral nervous system meet remain understudied. Combining single-cell transcriptomics, fate mapping, and parabiosis experiments, we show that five types of myeloid cells including three tissue-resident macrophage subpopulations, coexist in the mouse cochlea. The three macrophage subsets showed different potential functions in relationship with their specific topography across cochlear compartments. Further analysis revealed that they were partially derived from yolk sac progenitors during development, while in adulthood, most cochlear macrophages were long-term resident. Finally, we showed that cochlear macrophage morphology and density changed during aging. Our findings show that cochlea is a microenvironment with a unique heterogeneity of macrophages in terms of gene expression, spatial distribution, ontogeny, and function.

## Introduction

Tissue-resident macrophages are innate immune cells present in all tissues of the body, and they are involved in a wide range of functions depending on their anatomical location and organ-specific function ^1–3^. Despite having this wide range of functions, until recently, tissue-resident macrophages were considered a homogeneous population of cells across or within organs. In the past decade, however, fate-mapping studies and systems biology approaches, such as single-cell RNA-sequencing (scRNA-seq) analysis, have revealed a striking degree of heterogeneity related to ontogeny, transcriptional profiles, and self-renewal capabilities in macrophage populations within the same organ, suggesting microenvironment-specific phenotypes and functions ^1,2^.

Little is known about the identity of tissue-resident macrophages in the cochlea, a sensory organ sheltered by the blood-labyrinth barrier. While the cochlea was long thought to be immunologically privileged because of this barrier ^4^, more recent studies have revealed the presence of immune cells, mainly macrophages, in different compartments of the mouse and human inner ear ^5–7^. Cochlea as an organ on the border of central and peripheral nervous system represent a unique microenvironmental niche for macrophages. However, potential macrophage subpopulations in the cochlea are yet to be identified, and their ontogeny remain largely unknown. This lack of knowledge is an obstacle to a clear understanding of the connection between cochlear function and hearing impairments that could be caused by factors associated with the immune system, such as viral infections or aging.

Correlative immunofluorescent analyses of macrophages in *Csf1r* knock-out mice have suggested that, during development, cochlear macrophages have a dual origin. One population appears to originate from the yolk sac’s colony-stimulating-factor-1 receptor (CSF1R) –dependent erythro-myeloid precursors (EMP), mainly located in the spiral ganglion and the spiral lamina. Another population, independent from CSF1R signaling, likely originates from the fetal liver ^8^. In addition, cochlear macrophages of adult mice proposed to be maintained by bone marrow-derived immune cells, according to studies using chimeric mice obtained by bone marrow transplantation of lethally irradiated recipients, suggesting that circulating monocytes also contribute to the tissue-resident macrophage population ^9,10^. We and others, however, have shown that irradiation and injection of bone marrow precursors into the bloodstream are two confounding factors that cause an artificial infiltration of macrophages from peripheral blood to the central nervous system (CNS) and the eye ^11–13^. A similar effect may happen to the blood-labyrinth barrier of the cochlea. Using more physiological approaches such as fate-mapping or a parabiosis paradigm is therefore necessary to clarify the origin of cochlear macrophages.

In this study, we used unbiased scRNA-seq, genetic fate-mapping, and parabiosis experiments to understand the ontogeny, gene expression, spatial distribution—as well as the potential functions—of cochlear macrophages. We here provide the first mapping of cochlear myeloid cells, demonstrating that the immunogenic environment of this organ appears to be highly diverse and contains subsets analogous to several other tissues. Further, we revealed the properties, markers, and biological functionalities of each distinct subset, which establish their ontology. Our data indicate that the cochlea uniquely contains five distinct myeloid cell subpopulations, with core gene signatures similar to CNS microglia (brain and retina), peripheral nerve macrophages (sciatic nerve), and a peripheral immune organ (spleen). In addition, we uncovered evidences of unique gene signatures for each population, associated with their role in cochlea indicating the strong effect of cochlea microenvironment in imprinting the macrophage transcriptional program. By lineage-tracing the ontogeny of tissue-resident macrophages in development, adulthood, and aging, we demonstrated that cochlear myeloid cells partially originate from the yolk sac, with increased contribution from blood in a compartment-specific manner. This study, for the first time, provides the identity and diversity of cochlear myeloid cells, based on their origin and phenotypes in homeostasis and aging, and may have profound implications for discovering therapeutic targets for hearing loss.

## Results

### Single-cell RNA-sequencing revealed five myeloid cell subpopulations in the cochlea with core transcriptional differences

Few scRNA-seq approaches, have been applied to study the cochlea, and none with a focus on cochlea myeloid cells ^14–16^. As such, to generate an in-depth picture of the cochlea immune landscape we performed scRNA-seq to profile cochlear myeloid cells and their transcriptomic signatures in adult (6-week-old) wild-type (WT) C57BL/6J mice. We established a protocol to sort and enrich myeloid cells specifically from the cochlea by fluorescence-activated cell sorting (FACS), using the markers CD45, CD11b and F4/80 (**Figure S1A**). A total of 2,182 single-cell transcriptomes were subjected to quality control (see Materials and Methods), underwent unsupervised clustering, and were displayed using Uniform Manifold Approximation and Projection (UMAP). Five discrete myeloid cell clusters exhibiting distinct gene expression characteristics were identified within the cochlea (**Figure 1A**). We termed five cochlea-associated myeloid cell populations clusters: alpha (α), beta (β), gamma (γ), delta (δ), and epsilon (ε).

**Fig. 1.**
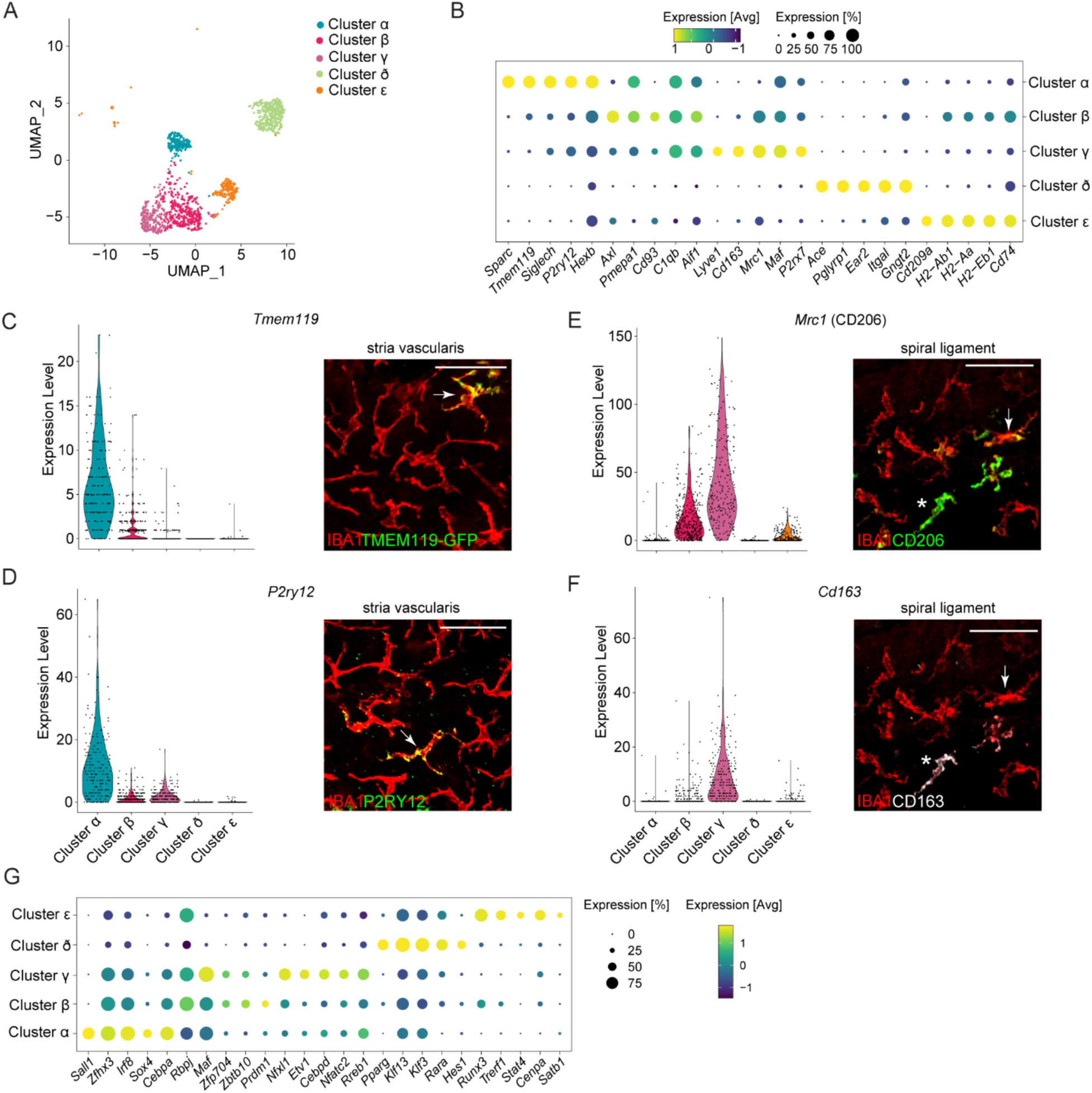
Single-cell RNA-seq of cochlear immune cells reveals unique transcriptomic heterogeneity. **A**. UMAP plot depicting five distinct clusters (cluster α to ε); n= 2,182 cells from 4 independent experiments. Each experiment contained cells from seven 6-week-old C57BL/6J mice. **B.** Dot plot of scaled expression of the top five markers for each cluster. Dot colors indicate the gene averaged expression, and dot sizes show the percentage of expression in the cluster. **C.** Violin plot of *Tmem119* gene expression in the cochlea clusters, showing high expression in the cluster α and representative confocal image of a whole mount preparation of the stria vascularis showing macrophages stained with IBA1 (red) and *Tmem119*-GFP (green) from the cluster α. **D.** Violin plot of *P2ry12* gene expression in the cochlea clusters showing high expression of *P2ry12* in the cells from the cluster α. Representative confocal image of the stria vascularis from a C57BL/6J mouse showing macrophages stained for IBA1 (red) and P2RY12 (green) macrophages from the cluster α. **E.** Violin plot of *Mrc1* (encoding for CD206) gene expression in the cochlea clusters showing high expression by clusters β and *γ*. Representative confocal image of spiral ligament from a C57BL/6J mouse showing macrophages stained for IBA1 (red) and CD206 (green) from the clusters β and *γ*. **F.** Violin plots of *Cd163* gene expression in cluster *γ but not cluster* β. Representative confocal image showing the same cells as in e. of a whole mount preparation of spiral ligament from a C57BL/6J mouse showing IBA1^+^ (red) and CD163^+^ (white) macrophages from cluster *γ*, to differentiate from cells in the *cluster* β being CD206^+^CD163^−^ (arrow) and the cluster *γ* being CD206^+^CD163^+^ (asterisk). **G.** Dot plot displaying the scaled expression of the top five transcription factors specifically upregulated in each cochlea cluster. Dot colors indicate the gene averaged expression, and dot sizes show the percentage of expression in the cluster.

Unique differentially expressed genes (DEGs) (log_2_FC > 1) between the cochlea myeloid cell clusters were used to initially distinguish possible cluster subtypes. Among the most highly expressed genes in cluster α (*Tmem119*, *P2ry12*, *Hexb*), cluster β (*Axl*, *Cd93*), and cluster γ (*Mrc1* (encoding CD206), *Cd163*, *P2rx7*) were genes known to be expressed by microglia or other macrophages, suggesting that they represented the macrophage populations of cochlea (**Figure 1B**) ^17^. In contrast, cluster ε expressed the C-type lectin gene *Cd209a* as well as antigen presentation-related genes such as *Cd74*, *H2-Ab1*, *H2-Aa* and *H2-Eb1* (**Figure 1B**), suggesting that this cluster was more similar to dendritic cells (DCs) ^18^. Cluster δ expressed unique transcripts compared with the other clusters, including *Ace*, *Ear2* and *Itgal*—genes typically expressed by non-classical monocytes in blood (**Figure 1B**), suggesting that this cluster was more similar to monocytes ^19^.

Next, we validated the presence of the three macrophage populations (α, β, γ) at the protein level in each compartment of the cochlea after dissection and immunostaining for corresponding markers (**Figure S2 and S3**). Cluster α cells were validated based on the co-expression of TMEM119 in *Tmem119*-GFP mice or P2RY12 with the pan-myeloid marker IBA1 in WT mice and were observed in the spiral lamina, the spiral ganglion, the stria vascularis, and the spiral ligament; but these cells were absent from the basilar membrane and the spiral limbus (**Figure 1C and 1D and Figure S2, S3 and S4**). Clusters β and γ were validated by staining for CD206 (encoded by *Mrc1*) and CD163. Based on our scRNA-seq data, *Mrc1* was expressed by both clusters β and γ, while *Cd163* could be used to distinguish between them since it was uniquely expressed by the cluster γ (**Figure 1E and 1F**). Staining for IBA1^+^ macrophages showed that some CD206^+^ macrophages expressed CD163, but all CD163^+^ macrophages expressed CD206, confirming that, at the protein level, we could distinguish both cluster β and γ subpopulations based on these markers (**Figure 1E and 1F and Figure S5**). Cluster β cells (CD206^+^CD163^−^) were detected in all compartments, with a strong presence in the stria vascularis, the spiral limbus, and the basilar membrane. In contrast, cluster γ cells (CD206^+^CD163^+^) were absent from the stria vascularis but present in other compartments.

Interestingly, the morphologies of macrophages were also highly diverse across all compartments. They represented a spectrum from ramified shape, often considered surveillant, in the stria vascularis to a more amoeboid shape, often considered fully activated cells that play a role in phagocytosis, in the spiral ligament (**Figure 1C-1F and Figure S4 and S5**).

To investigate the molecular determinants of the diverse populations of myeloid cells in the cochlea, we characterized the transcriptional regulators that were upregulated in each cochlea-associated myeloid population by performing differential expression analysis on known transcription factors (log_2_FC threshold > 0.4, adjusted p-value < 0.05). The five most highly expressed transcription regulators present in each subpopulation confirmed cluster-specific expression of the individual transcription factors, except for a few overlaps for clusters β and γ (**Figure 1G**). Genes encoding the transcription factors *Sall1*, *Cebpa*, *Irf8*, *Zfhx3* and *Sox4* were specifically expressed in cluster α, *Etv1*and *Cepd* in cluster γ, *Runx3* and *Stat4* in cluster ε, and *Rara* and *Pparg* in cluster δ (**Figure 1G**). Both cluster β and γ were enriched in *Rbpj* and *Maf*, which are master regulators of macrophage polarization and differentiation, respectively (**Figure 1G**) ^20,21^.

Collectively, the heterogenous gene expression at the single-cell level, together with the spatial distribution of the corresponding myeloid-specific populations across compartments, suggest a previously unappreciated heterogeneity across cochlear macrophages.

### Myeloid cells subpopulations in the cochlea exhibited strong transcriptional similarity with macrophages across the nervous system and immune organs

Emerging evidence suggests that different populations of tissue-resident macrophages have strong degrees of organ-specific transcriptional identity that are shaped by their local environments and unique functions ^1,2^.

To further define the identity of the five myeloid subpopulations that we found in the cochlea, we compared their transcriptomic profile to well-characterized, tissue-resident macrophages from seven different organs (**Figure 2A**). We isolated myeloid cells from a secondary lymphatic organ rich in immune cells (spleen), a peripheral immune compartment (peritoneal cavity), a respiratory organ (lungs), a metabolic organ (liver), different parts of the CNS (retina, brain), and a part of the peripheral nervous system (sciatic nerve) and compared them to cochlear myeloid cells (**Figure S1**). The myeloid cells from these organs represent a wide spectrum of tissue-specific transcriptomes and functions.

**Fig. 2.**
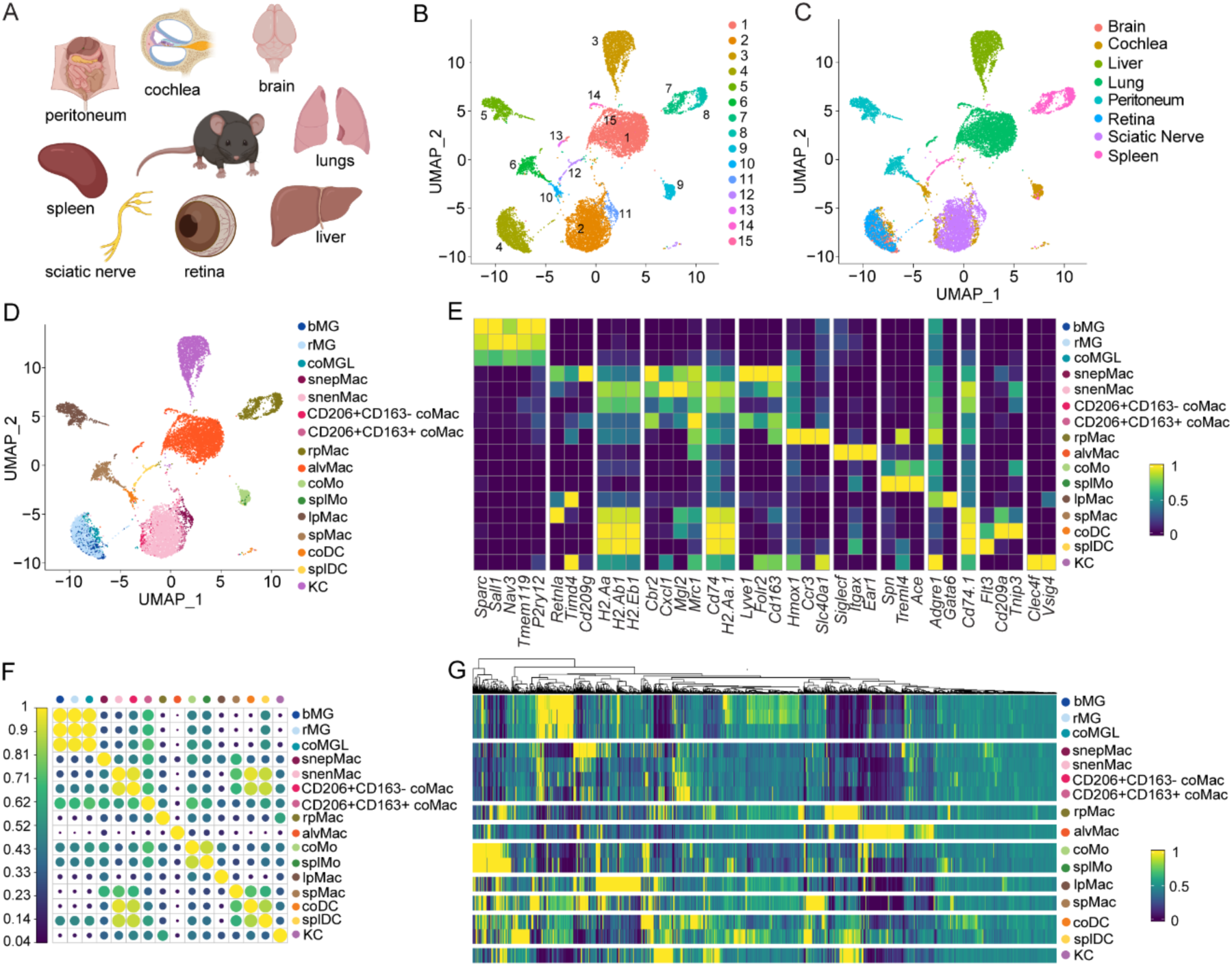
Single-cell RNA profiling of myeloid cells in multiple organs including the cochlea. **A.** In addition to cochlear macrophages, macrophages from the brain, lungs, liver, retina, sciatic nerve, spleen, and peritoneum in steady-state conditions were sorted and processed using 10x Genomics Chromium Next GEM Single-cell protocol (see gating strategy in fig. S1). **B.** UMAP plot of all macrophages from all tissues showing 15 different clusters; n = 14,662 cells, from 4 independent experiments (cochlea) or 3 independent experiments (all other tissues). Each experiment contained cells from three 6-week-old C57BL/6J mice (brain, liver, peritoneal macrophages, lung, spleen) or seven 6-week-old C57BL/6J mice (cochlea, retina, sciatic nerve). **C.** UMAP plot of all macrophages in which each cluster was identified according to its tissue of origin. Each color is assigned to a different tissue of origin. **D.** UMAP plot of all macrophages in which cell type identities were assigned to the clusters based on differentially expressed markers identified in the literature. Different color indicate the assigned cellular identity, respectively. **E.** Heatmap depicting the average expression of different marker genes for each cluster known from current literature. **F.** Correlation dot plot comparing the different cell clusters with each other and indicating relationships based on the top 1811 differentially expressed genes. **G.** Heatmap depicting the average expression of the top 2000 most highly variable features that will be used in downstream analysis.

To minimize experimental artifacts such as cell activation due to enzymatic digestion that is necessary for processing several tissues such as the liver, cochlea, and sciatic nerve, we implemented a systematic correction of the gene signatures associated with and genes highly correlated with enzymatic digestion (see Material and Methods for the quality control process)^22,23^.

After quality control, a total of 14,662 single-cell transcriptomes with 3,730 median genes per cell from all eight organs, or parts thereof, were subjected to unsupervised clustering and UMAP for dimension reduction, resulting in 15 transcriptionally distinct clusters in our combined analysis (**Figure 2B**). Upon identification of the tissue of origin for each cell, UMAP and Louvain clustering algorithms revealed a clear separation among most of the tissues (liver, lung, spleen, peritoneum, and sciatic nerve), except for brain and retinal microglial cells that as expected clustered together ^13,24^ confirming organ-specific macrophage transcriptomes (**Figure 2C**).

Strikingly, in contrast to all the other tissues, cells isolated from the cochlea did not cluster in an organ-specific pattern but rather divided into five subclusters, each of which was closely related to other tissues (**Figure 2C**). One cochlea subpopulation (cluster α) clustered close to brain and retinal microglial cells, two subpopulations (clusters β and γ) associated with sciatic nerve macrophages, and two subpopulations (clusters δ and ε) clustered with splenic cells (**Figure 2C**). None of the cochlear myeloid cells, in contrast, clustered with liver, lung or peritoneal macrophages. Based on known markers, we confirmed the cellular identities in each tissue (**Figure 2D and 2E**), and named each cochlea cluster based on their proximity to other tissue myeloid cell populations and their expression of similar markers. We named the cochlea cluster α close to brain and retinal microglial cells “cochlea microglia-like cluster (coMGL)” and identified expression of signature genes known for microglia including *P2ry12*, *Tmem119*, *Sparc*, *Sall1*, and *Nav3*, albeit to a lesser extent than brain and retina microglial cells (**Figure 2D and 2E**) ^13,25^. We named cochlea subpopulations clustering with sciatic nerve macrophages “cochlea sciatic nerve-like macrophages”; we further differentiated these based on their gene and protein expression of *Cd163* and *Cd206* and termed them CD206^+^CD163^−^ coMac (cluster β) and CD206^+^CD163^+^ coMac (cluster γ) (**Figure 1D and 1E**). Interestingly, these two clusters showed some similarities to previously identified sciatic nerve endoneurial and epineurial macrophage subtypes (snendoMac, snepiMac) expressing *Cd74* and *Lyve1*, respectively ^22^. Finally, we named the cochlea subgroup clustering with spleen monocytes, which also expressed *Spn*, *Treml4* and *Ace*, albeit at a lower level, “cochlear monocytes (coMo)” (cluster δ), and we named the cochlea cluster close to spleen DCs “cochlear DC (coDC)” which expressed *Flt3*, *Cd209a* and *Tinp3* (cluster ε). (**Figure 1D and 1E**).

We further interrogated the level of transcriptional similarity between these cochlea populations and all other populations using Pearson correlation calculated based on the top 1811 most variable genes (2000 most variable genes minus the enzymatic dissociation-associated genes). The CoMGL subpopulation shared a very strong correlation with brain and retinal microglia (0.99). The CD206^+^CD163^−^ coMac cluster shared a robust correlation with snendoMac (0.96) of sciatic nerve, while CD206^+^CD163^+^ coMac cluster was slightly more correlated to snepiMac (0.66) than to snendoMac of sciatic nerve (0.56) (**Figure 2F**). The CoMo shared a very strong correlation with spleen monocytes (0.93) and coDC with spleen DCs (0.87) (**Figure 2F**). The CD206^+^CD163^−^ coMac shared a strong correlation with coDC (0.92), which was surprising. This may have been due to the high level of antigen presentation-related genes in both groups (**Figure 2E and 2F**). These close relationships between each cochlea clusters and their proximate tissue was confirmed by the parallel transcriptional signature they shared (**Figure 2G**). Overall, the correlation was lower between the cochlea clusters than between cochlea clusters and their proximate non-cochlea tissue.

To obtain a better understanding of their functions, we investigated the functional profiles of the differentially expressed genes among all cochlea clusters with gene ontology (GO)–term analysis. CoMGL-enriched genes were related to “cytoskeleton organization” and glial cell processes including “gliogenesis”, “glial cell migration” and “synapse pruning” (**Figure 3A and 3B** and **Figure S6**), confirming their functional similarities to microglia. Genes related to “T-cell differentiation”, “T-cell proliferation”, and “T-cell activation” functions were expressed by all clusters, although they were predominantly enriched in the coMo and coDC clusters. Functions in relation to antigen presentation and MHC class II complex, however, were almost exclusively linked to the coDC and, to a lesser extent, CD206^+^CD163^−^ coMac cluster (**Figure 3A and 3B** and **Figure S6**). In addition, the CD206^+^CD163^−^ coMac cluster expressed TNF genes and interleukin-6 production-related genes. These genes are related to various classical macrophage functions such as the inflammatory response^26,27^. This cluster also expressed genes involved in cell adhesion and myeloid cell differentiation (*Rbpj*, *Cdk6*, and *Id2*) (**Figure 3A and 3B** and **Figure S6**).

**Fig. 3.**
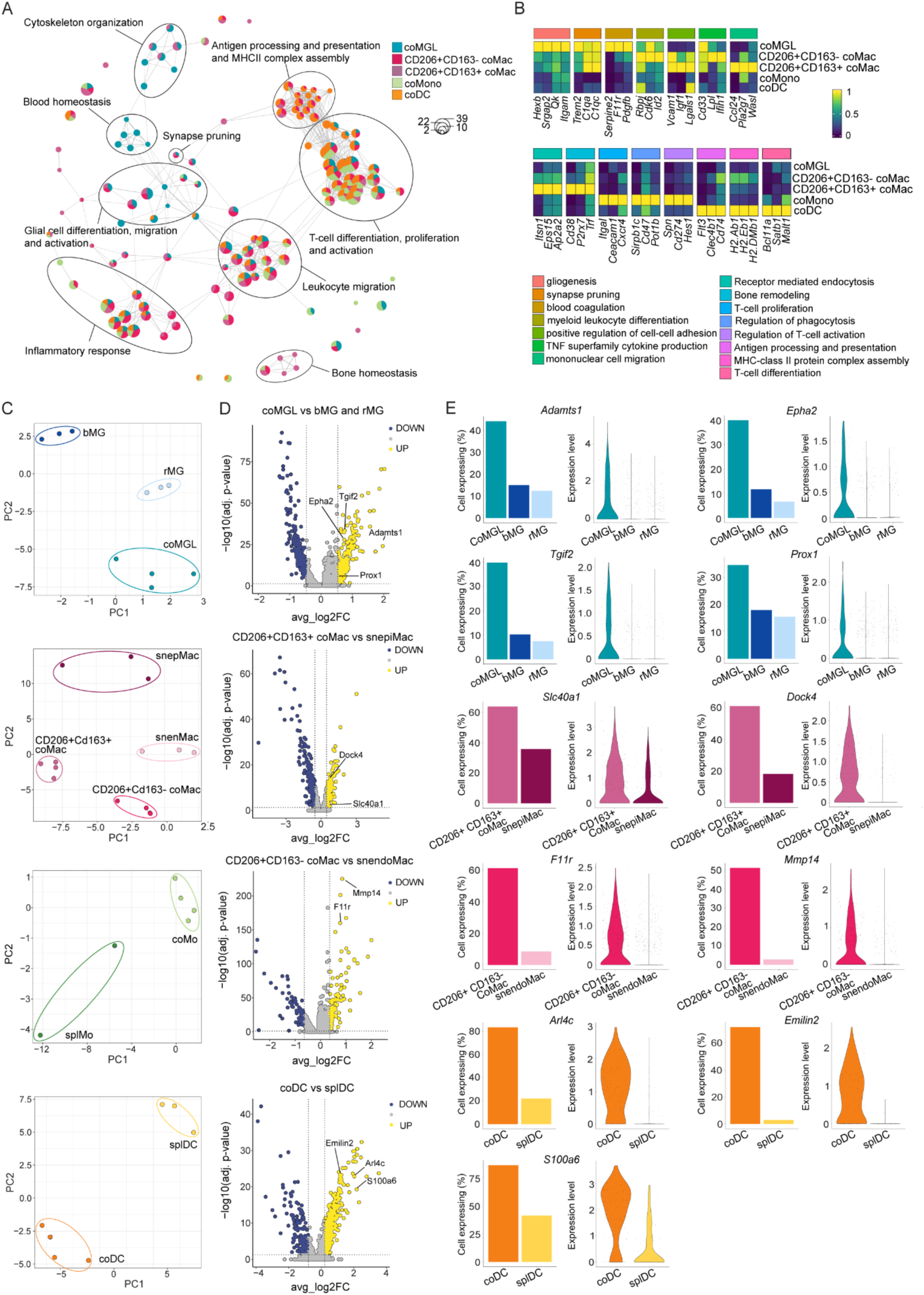
Transcriptional analysis reveals specific functions and gene expression of cochlear macrophages. **A.** Gene ontology (GO) network based on genes differentially expressed between cochlea clusters. Each pie chart represents a GO term that displays the relative number of term-related genes upregulated in each cochlea cluster. Lines connect GO terms with overlapping genes. GO terms referring to related functions are grouped and circled together. **B.** Heatmap depicting selected gene average expressions from selected GO terms highly represented in one of the five cochlea clusters (p-filtered; scaled). **C.** PCA analysis of each cochlea cluster with transcriptionally close clusters from other tissues. **D.** Volcano plot depicting differential expression of selected genes between each cochlea cluster including coMGL, CD206^+^CD163^+^ and CD206^+^CD163^−^ coMac, and coDC with their transcriptionally related clusters from other tissues. Yellow and blue dots display upregulated and downregulated genes, respectively. **E.** Bar graph showing the percent of expression and violin plot depicting the level of expression of genes from the comparison in (C) between a specific cochlea cluster and its analog cluster in other tissues. Genes were selected based on their known biological relevance for inner ear biology.

Some of the cochlea clusters—in addition to exhibiting common functional features with analogue clusters in other tissues—exhibited key functional differences with their analog cluster. In coMGL cells, some of the most-expressed differentially regulated genes, as compared to brain and retina microglia, were related to blood homeostasis, including platelet activation (*Serpin2*, *F11r* and *Pdgfb*), angiogenesis, coagulation, and hemostasis (**Figure 3A and 3B** and **Figure S6**). Whereas CD206^+^CD163^−^ and CD206^+^CD163^+^ coMac clusters, which are similar to sciatic nerve macrophages, were enriched in genes related to complement activation important for synapse pruning and usually absent in sciatic nerve macrophages (**Figure 3A and 3B** and **Figure S6**). Interestingly, CD206^+^CD163^−^ and CD206^+^CD163^+^ coMac subsets were present in the basilar membrane of the cochlea where the hair cell – spiral ganglion synapses reside while coMGL were absent, suggesting that they might be exclusively responsible for synapse pruning (**Figure S4 and S5**). CD206^+^CD163^+^ coMac cluster cells also displayed specific enrichment of genes linked to bone homeostasis, such as *Cd38*, *Trf* or *P2rx7* (**Figure 3A and 3B** and **Figure S6**). While bone metabolism genes could be the result of sample contamination from bone pieces from the cochlea capsule, none of the classical markers of osteoclast (Acp5, Ctsk and Nfatc1) were expressed by the CD206+CD163+ coMac population.

We hypothesized that, along with their core gene profiles, each macrophage population has a unique gene signature most likely associated with their role in the tissue. PCA analysis of each cochlea cluster and their analog clusters from other tissues revealed that, despite very high correlation (**Figure 2F**), PC1 and PC2 were driven by the tissue of origin, suggesting strong imprinting of the environment to the transcriptional signature (**Figure 3C**). To ascertain a unique identity for each cochlea cluster, we looked for any genes unique to each cluster compared to their analog cluster in other tissues (**Figure 3D**). In coMGL cells, we found that *Adamts1*, *Epha2*, *Tgif2*, and *Prox1* were overexpressed compared to brain and retinal microglia (**Figure 3D and 3E**). The CD206^+^CD163^+^ coMac cluster uniquely expressed *Slc40a1* and *Dock4*, compared to sciatic nerve epineurial macrophage subtypes (**Figure 3D and 3E**). The CD206^+^CD163^−^ coMac cluster uniquely expressed *F11r* and *Mmp14*, compared with the sciatic nerve endoneurial macrophage subset, and coDC uniquely expressed *Emilin2*, *S100a6* and *Arl4c,* compared with splenic DCs (**Figure 3D and 3E**).

Overall, our data depicts a strong heterogeneity of myeloid cells in the cochlea and place the cochlea at the frontier between the central nervous system, the peripheral nervous system, and the periphery.

### Lineage tracing revealed ontogeny and self-renewal heterogeneity in cochlear macrophages

Other potential sources of heterogeneity among cochlear macrophages are the relative contributions from distinct lineages and the mechanism of tissue-resident macrophage replenishment throughout the life of the animal. While microglia in the brain are entirely derived from precursors in the yolk sac early during development, sciatic nerve macrophages are primarily derived from fetal liver progenitors ^22,28,29^. Similarly, microglia maintenance is driven entirely through self-renewal in the adult independent of circulating monocytes, while sciatic nerve macrophages are replaced gradually by circulating monocytes after birth ^22,28,30^. As cochlea myeloid cells contain both microglia-like and sciatic nerve-like macrophages, we next sought to unravel the ontogeny of cochlear tissue-resident macrophages using fate-mapping models.

To label early embryonic macrophage precursors in the yolk sac, we generated *Cx3cr1*^CreERT2/+^:*Rosa26^YFP/YFP^*mice and pulse-labeled *Cx3cr1*-expressing cells with 4-OH-tamoxifen (4-OH-TAM) at E9.5 (**Figure 4A** and **Figure S7A**). The *Cx3cr1*^CreER-T2^ line has been reported to show TAM-independent recombination events; therefore, we compared mice that received 4-OH-TAM injections to untreated mice of the same genotype (**Figure 4B**)^13,31^. Brain microglia from age-matched littermates with the same genotype were analyzed as references for the recombination efficiency. In comparison to the brain (36.2 ± 7.5 % of IBA1^+^ expressing YFP), cochlea myeloid cells showed low YFP labeling (around 10%), suggesting that the fetal liver may be an additional source of cochlear macrophages during development (**Figure 4B** and **Figure S7B**) ^22,32^. Macrophages in the basilar membrane, spiral lamina, spiral ganglion, stria vascularis and spiral ligament showed significantly higher percentages of YFP^+^ cells in the IBA1^+^ population, compared to untreated mice; while the percentage of YFP^+^ macrophages in the spiral limbus was not significantly different from untreated mice (**Figure 4B**).

**Fig. 4.**
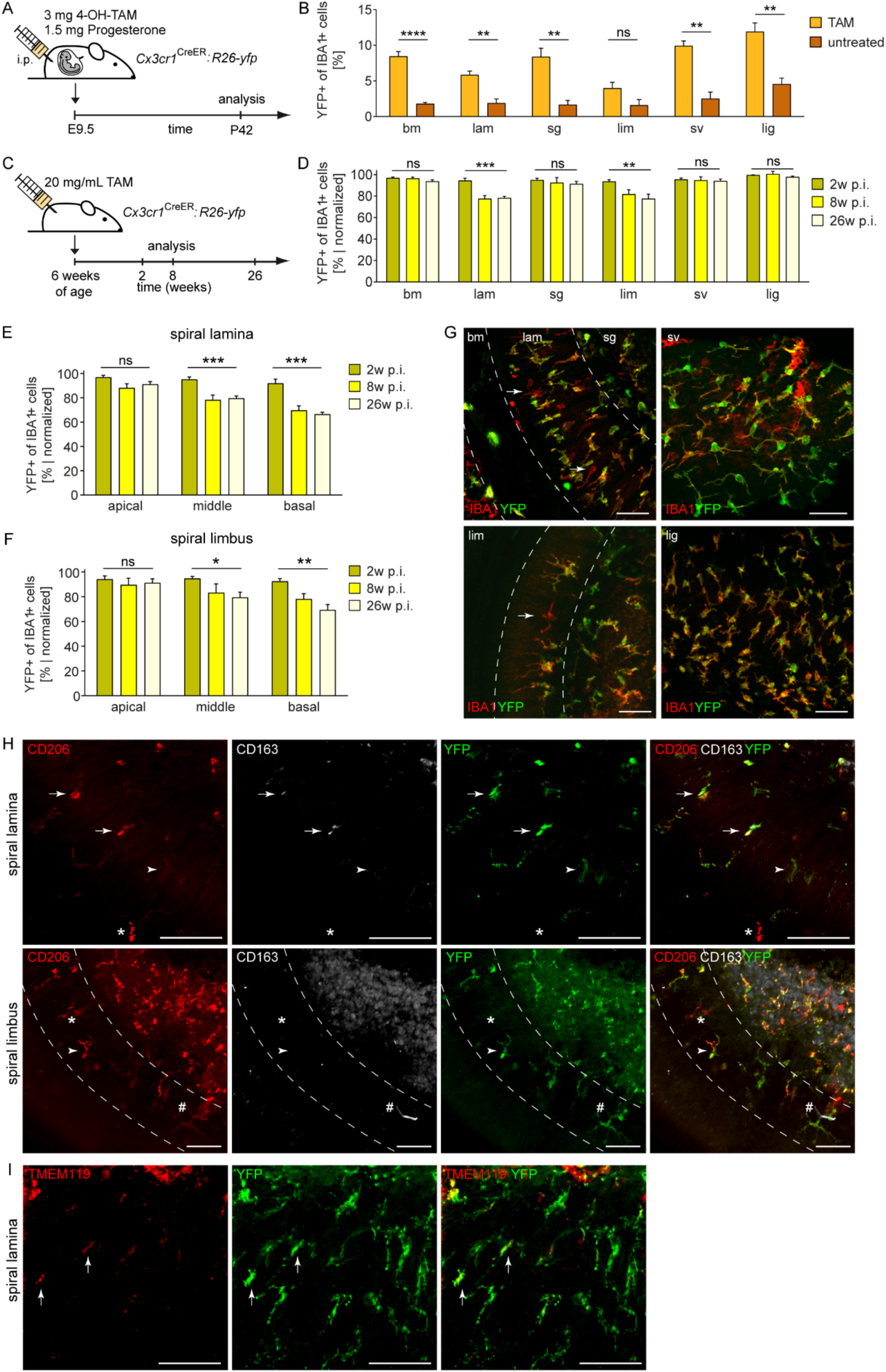
Dynamics and ontogeny of cochlear macrophages in steady-state conditions. **A.** Scheme of a fate-mapping experiment using *Cx3cr1^CreERT^*^2^*:Rosa26-YFP* mice. 4-Hydroxy-Tamoxifen (4-OH-TAM) and progesterone injections were performed at embryonic day 9.5 (E9.5). Mice were evaluated at postnatal day 42 (P42). **B.** Quantitative analysis of YFP expression in IBA1^+^ macrophages in the different cochlea compartments: basilar membrane (bm), spiral lamina (lam), spiral ganglion (sg), spiral limbus (lim), stria vascularis (sv), and spiral ligament (lig) of TAM-induced and untreated *Cx3cr1^CreERT^*^2^*:Rosa26-YFP* mice at P42. Bars represent means ± s.e.m; n=7 (TAM) and n=3 (untreated) animals. ** p < 0.01 and **** p < 0.0001. **C.** Scheme of turnover experiment using adult *Cx3cr1^CreERT^*^2^*:Rosa26-YFP* mice. Tamoxifen (TAM) injection was performed on postnatal day 42 (P42). Mice were evaluated at 2-, 8– and 26– weeks post-injections. **D.** Percentage of YFP labeling in IBA1^+^ macrophages of the different cochlea compartments: basilar membrane (bm), spiral lamina (lam), spiral ganglion (sg), spiral limbus (lim), stria vascularis (sv), and spiral ligament (lig). Bars represent means ± s.e.m; n = 11 (2 and 26 weeks p.i) and n = 6 (8 weeks p.i.), ** p < 0.01 and *** p < 0.001. **E-F.** Percentages of YFP labeling in IBA1^+^ macrophages of the spiral lamina (e) and the spiral limbus (f). Bars represent means ± s.e.m; n = 11 (2 and 26 weeks p.i) and n = 6 (8 weeks p.i.), * p < 0.05, ** p < 0.01, and *** p < 0.001. **G.** Representative immunofluorescence images from the different cochlear compartments from whole mounts of *Cx3cr1^CreERT^*^2^*:Rosa26-YFP* mice at 26 weeks p.i. Short-lived IBA1^+^ YFP^−^ are shown with arrows. Scale bars = 50 μm. **H.** *Cx3cr1^CreERT^*^2^*:Rosa26-YFP* mouse cochlea shows long-lived YFP^+^ (green) CD206^+^ (red) CD163^+^ (white) (arrows) and YFP^+^CD206^+^CD163^−^ (arrowhead) and short-lived YFP^−^ CD206^+^CD163^+^ (hash) and YFP^−^CD206^+^CD163^−^ (star) coMacs. Pictures are representative of two animals. Scale bars: 50 μm. **I.** Confocal image of the spiral lamina from a *Cx3cr1^CreERT^*^2^*:Rosa26-YFP* mouse cochlea. TMEM119^+^ (red) YFP^+^ (green) coMGL (arrows) indicating long-lived character. Pictures are representative of two animals. Scale bars = 50 μm.

Previous studies using chimeric mice obtained by bone marrow transplantation of lethally irradiated recipients have suggested that bone marrow–derived monocytes continually replace cochlear macrophages ^9,10^. Yet this approach has been shown to suffer from experimental confounds, because irradiation, which promotes the disruption of the blood-brain barrier, and the injection of hematopoietic stem cells into the circulation are both thought to induce artifactual infiltration into the tissue ^11,12^. We therefore took advantage of our conditional *Cx3cr1*^CreER-T2^:*Rosa26-YFP* reporter mouse model to track macrophages over time to study the maintenance of cochlea tissue-resident macrophages. To identify bone marrow-derived circulating monocytes, we treated 6-week-old *Cx3cr1*^CreER-T2^:*Rosa26*-YFP reporter mice with TAM and analyzed cochlea 2-, 8– and 26– weeks after TAM injections (**Figure 4C**).

Recombination efficiency was confirmed by FACS analysis of blood and brain (**Figure S8A and S8B**). FACS analysis of blood revealed YFP expression in 41.01 ± 2.32% of monocytes at 1-week post-TAM administration (p.i.), as previously published, which rapidly decreased to an undetectable level at 8 and 26 weeks p.i. (**Figure S8A**) ^33^. Brain myeloid cells, including microglia and border-associated macrophages, in contrast, reached high levels of expression as soon as 2 weeks p.i. (89.8 ± 2.3%), which increased to 96.8 ± 1.2% of YFP^+^ brain myeloid cells at 26 weeks p.i, allowing us to use YFP percent in the brain to individually normalize YFP expression in the cochlea. (**Figure S8B**).

Histological analysis of the cochlea showed a high percentage of YFP^+^ macrophages in all compartments at 2 weeks p.i. (**Figure 4D**). Percentages of YFP^+^ macrophages were not different at any of the time points assessed in the basilar membrane, spiral ganglion, stria vascularis, and spiral ligament with over 90% of IBA1^+^ macrophages YFP^+^ at any given time (**Figure 4D**). In contrast, we observed that the fraction of YFP^+^ macrophages in the spiral lamina and the spiral limbus decreased to 76.03 ± 2.2% and 76.05 ± 4.7%, respectively, at 26 weeks p.i. (**Figure 4D**). These findings show that, while tissue-resident cochlear macrophages represented a stable population in most compartments, a slow and progressive contribution of circulating monocytes to the macrophage pool occurs in the spiral lamina and spiral limbus.

Cochlear macrophages may not only differ between compartments but also within the same compartment, dependent on the turn of the cochlea, which adds another level of complexity. Therefore, we subdivided and investigated each compartment in the basal, middle, and apical turns (**Figure 4E to 4G** and **Figure S3 and S9**). YFP^+^ expression in macrophages of the basilar membrane, spiral ganglion, stria vascularis and spiral ligament remained stable over time, regardless of the turn (**Figure S9**). Our detailed analysis, however, revealed that the decrease of YFP^+^ macrophages previously detected in the spiral lamina and the spiral limbus was observed only in the basal and the middle turns (**Figure 4E to 4G** and **Figure S9**). These findings reveal that apical cochlear turns lacked significant monocyte contribution and contained a stable macrophage population—even in compartments experiencing monocyte replenishment (**Figure 4E to 4G** and **Figure S9**).

The turnover of tissue-resident macrophages can be investigated either by labeling the macrophage itself, as described above, or by doing the inverse and labeling monocytes in the peripheral blood to identify the monocyte-derived macrophages in the tissue ^11,34^. To confirm our previous results, we performed parabiosis to establish a shared circulation between *Ubiqutin*-GFP (*Ubc*-GFP) and C57BL/6J mice; then, we analyzed the contribution of GFP^+^ monocytes to the pool of IBA1^+^ cochlear macrophages in the C57BL/6J mouse at 4– and 28-weeks post-surgery (p.s.) (**Figure S10A**). FACS analysis of blood at 2 weeks post-surgery (p.s.), showed efficient blood sharing as previously reported (**Figure S11A**) ^11,34^. Additional analysis of the blood and the spleen at 4 and 28 weeks p.s. confirmed strong labeling of blood– and spleen-derived monocytes, T cells, and B cells (**Figure S11A to S11D**), confirming successful chimerism. In contrast, but as expected, GFP^+^ cells were not detectable in the brain of C57BL/6J *Ubc*^+/+^ partner mice (**Figure S11E**). Histological analysis of the cochlea revealed, in alignment with the results obtained with the *Cx3cr1*^CreER-T2^*:Rosa26-YFP* model, GFP^+^ cells were only found in the spiral lamina and spiral ligament of the C57BL/6J mouse, while all other compartments remained devoid of any detectable GFP^+^IBA1^+^ macrophages (**Figure S10A to S10F**). We again subdivided our analysis by cochlear turns, namely basal, middle, and apical (**Figure S3**). We found that GFP^+^IBA1^+^ macrophages could be detected in the basal and middle turns of the spiral lamina and spiral limbus but were absent from the apical turn in both compartments, comprehensively revealing the same pattern as in the *Cx3cr1*^CreER-T2^*:Rosa26-YFP* mice (**Figure S10G to S10N**).

We next assessed the potential contribution of infiltrating monocytes to the identified cochlea macrophage subpopulations by co-immunostaining for TMEM119, CD206 or CD163 with the YFP expressed by resident macrophages (**Figure 4H and 4I**). All TMEM119^+^ coMGL cells were labeled with YFP, suggesting that local self-renewal rather than blood monocyte replacement was responsible for the maintenance of this population. However, a portion of CD206^+^CD163^−^ and CD206^+^CD163^+^ coMac cells did not express YFP, indicating a partial replacement by blood monocytes for these two populations (**Figure 4H and 4I**).

Collectively, this data suggests ontogeny and origin of macrophages during development and in adulthood also contributes to the diversity of cochlea macrophage populations.

### Cochlea macrophage dynamics and phenotypes adapt to aging in C57BL/6J mice

Aging is known to influence macrophage dynamics and phenotypes, which in turn impacts key physiological functions for tissue maintenance and repair. In that regard, emerging evidence supports the hypothesis that macrophages may contribute to age-related hearing loss, affecting around 30% of people over 65 years old, through complex interactions with other cell types ^35^. C57BL/6J mice are known to experience early-onset, age-related hearing loss due to an inbred mutation in the *Cdh23* gene, which affects cochlear hair cell survival ^36^.

We thus sought to investigate changes in our previously identified macrophage subsets in the cochlea of C57BL/6J mice during aging. Changes in macrophage morphology during aging have previously been reported in other studies in mice and humans ^37–39^. In 70-week-old mice, IBA1^+^ macrophages appeared to be larger and with a more rounded morphology, an appearance that is associated with less ramification, specifically in the basilar membrane and the spiral lamina (**Figure 5A)**. Similar to previous studies, different macrophage morphologies along the basilar membrane were also present in the young mouse, with macrophages in the basal turn displaying a round and amoeboid shape, compared with the ramified filopodia morphology in the apical turn, as a sign of an age-associated change in appearance of macrophage with potential functional alterations (**Figure 5A**) ^5,37–39^.

**Fig. 5.**
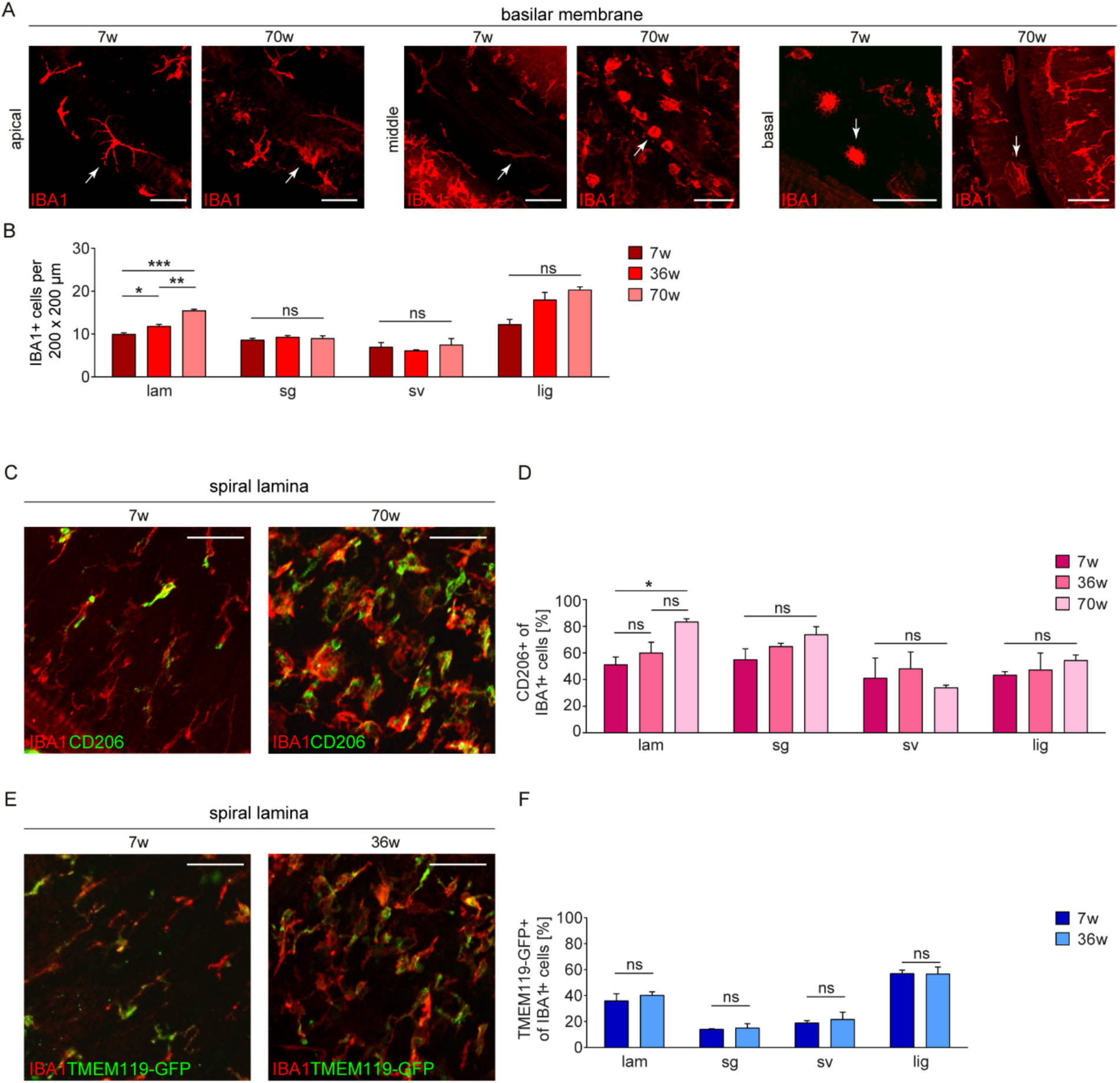
Cochlea macrophage dynamics and phenotypes change following early onset hearing loss in C57BL/6J mice. **A.** Representative images of the apical, middle and basal turns of the basilar membrane showing IBA1+ macrophages (in red – white arrows) in 7– and 70-weeks-old C57BL/6J mice. Scale bars = 50 μm. **B.** Quantitative analysis of IBA1^+^ macrophage density (cell counts in two fields of 200μm x 200μm per animal) in the spiral lamina (lam), spiral ganglion (sg), stria vascularis (sv), and the spiral ligament (lig) of 7-, 36-, and 70-weeks-old C57BL/6J mice. All quantifications were done in the middle turn of the cochlea. Bars represent means ± s.e.m.; n=3 mice per time point, * p < 0.05, ** p < 0.01, and *** p < 0.001. Of note, macrophage density was too low in the basilar membrane and the spiral limbus to be accurately quantified. **C.** Representative images of the spiral lamina stained with IBA1 (in red) and CD206 (in green) depicting CD206+ coMacs in 7– and 70-weeks-old C57BL/6J mice. Scale bars=50μm. **D.** Percentage of IBA1^+^ macrophages expressing coMac marker CD206 in 7-, 36-, and 70-weeks-old C57BL/6J mice. Bars represent means ± s.e.m.; n=3 per time point, * p < 0.05. **E.** Representative images of the spiral lamina stained with IBA1 (in red) and TMEM119 (in green) depicting TMEM119^+^ coMGL in 7– and 26-weeks-old *Tmem119*-GFP mice (on a C57BL/6J background). Scale bars=50μm. **F.** Percentage of IBA1^+^ macrophages expressing coMGL marker TMEM119 in 7-, and 36-weeks-old *Tmem119*-GFP mice. Bars represent means ± s.e.m.; n=3 per time point.

Macrophage numbers dramatically increase in the cochlea, especially during the first postnatal week ^37^, but little is known about the dynamics of macrophages at later stages or during ageing. We found that IBA1^+^ macrophage densities did not increase in the spiral ganglion, the stria vascularis and the spiral ligament in older mice (**Figure 5B**). In contrast, we observed an increased density of IBA1^+^ macrophages over time in the spiral lamina (**Figure 5B**). While, spiral limbus and basilar membrane macrophage densities were not high enough to be accurately quantified.

We next sought to determine if this increase was specifically driven by TMEM119^+^ coMGL or sciatic nerve-like coMac subsets, which were observed to be primarily found within the six cochlear compartments. While we did not observe a change in the proportion of the coMGL subset, the CD206^+^ coMacs (including CD163^+^ and CD163^−^ macrophages) did increase in the spiral lamina (**Figure 5C to 5F**). These data suggest that the sciatic-nerve-like macrophage subset is the main driver of the macrophage increase in the spiral lamina over time (**Figure 5C to 5F**). Collectively, our data suggest that macrophages change due to ageing, as indicated by both their density and morphologies and a certain subset of macrophages in cochlea is responsible for the age-related changes.

## Discussion

The discovery of macrophages in cochlea dates back to 1988 ^40^ primarily based on simple histological analysis. Yet this analysis lacked features of molecular identity. Later studies, utilizing bone marrow-transplantation, suggested that cochlear macrophages originated from circulating monocytes ^9,10^. This basic framework explaining the ontogeny of cochlear macrophages has remained largely intact. Yet more recently, extensive fate-mapping studies and scRNA-seq approaches have revealed that even though macrophages are found as resident immune cells in all mammalian organs, macrophage subpopulations are specialized to meet the demands of each host tissue ^1^. Here, we reveal that the mouse cochlea harbors a unique heterogeneous population of macrophages. Our data uncovers the transcriptional diversity, ontogeny, self-renewal capabilities, and sub-tissular niches of cochlear macrophages.

Using scRNA-seq, we show that five independent populations of CD45^+^CD11b^+^ myeloid cells are present within the cochlea. Profiling cochlear myeloid cells against myeloid cells from seven well-characterized murine organs demonstrates that these five populations represent three subpopulations of tissue-resident macrophages, one monocyte population, and one DC population. The three macrophage subsets shared core transcriptional similarity with brain and retina microglial cells, and with sciatic nerve epineurial and endoneurial macrophages, while the monocyte and DC populations were similar to spleen monocytes and dendritic cells, respectively ^18,19,22^. We focused our analysis on the three macrophage subsets, because they resemble tissue-resident macrophages, and confirmed their identity and spatial distribution at the protein level. We named the population similar to brain and retina microglial cells coMGL (cochlea microglia-like cluster), the population similar to sciatic nerve epineurial macrophages CD206^+^CD163^+^ coMac, and the population similar to sciatic nerve endoneurial macrophages CD206^+^CD163^−^ coMac. This three-cell population model of the cochlea tissue-resident macrophages thus provides a framework for understanding their biology and function.

We leveraged the single-cell resolution of our data to correlate the potential role for each population in the context of cochlear anatomy and function. We hypothesized that, along with their core gene profiles, each macrophage population has a unique gene signature most likely associated with their role in the tissue. CoMGL highly expressed blood homeostasis-related genes compared to microglia in brain and retina. We propose that coMGL cells are responsible for the regulation of the vasculature and blood homeostasis. To support this hypothesis, aside from genes encoding transcription factors important for myeloid cells and microglial cell development and function (*Sall1*, *Cebpa* and *Irf8*), we found that coMGL express two transcription factors, *Zfhx3* and *Sox4*, previously associated with angiogenesis ^41,42^. Therefore, our data provide an insight into why coMGL cells reside in distinct and highly vascularized compartments of cochlea such as the stria vascularis and the spiral ganglion but were not found in the relatively less vascularized basilar membrane that is nurtured by the surrounding endolymphatic fluid. Similarly, unlike sciatic nerve macrophages, CD206^+^CD163^−^ coMac and CD206^+^CD163^+^ coMac expressed synapse pruning-related genes and were located in the basilar membrane and the spiral limbus near the hair cell – spiral ganglion synapses. In addition, despite the presence of DCs, CD206^+^CD163^−^ coMacs were highly enriched in MHC-II, T cell regulation and antigen processing-related genes, whereas CD206^+^CD163^+^ coMacs were enriched in genes important for fluid shear stress and bone metabolism, suggesting they have a unique tissue-specific role in the cochlea. These data highlight the key role that the tissue microenvironment plays in reprogramming macrophage populations to support their tissue-required function in homeostasis and disease.

Further, a number of genes were uniquely expressed by cochlear macrophages, suggesting these genes may be the niche-dependent signature of cochlea myeloid cells clusters. Strikingly, previous studies have shown many of these genes are linked to hearing loss (*Adamts1*, *Slc40a1*, *Epha2*, *Dock4* and *F11r*), or already described to be expressed (*Prox1, Emilin2*, *S100a6*, *Arl4c, and Mmp14*) in the cochlea ^43–52^. This is significant because it may allude to a role for macrophages in the etiology of a range of hearing loss conditions.

It has been established that several macrophage subpopulations exhibiting distinct gene siganture can co-exist within one tissue. For example, we have previously shown that in the CNS, microglial cells (Hexb^+^, Mrc1^−^, Lyve1^−^) co-exist with border-associated macrophages (Lyve1^+^, Mrc1^+^, Hexb^−^) ^3^. Interestingly this framework has recently been extended, as it was proposed that two or three macrophage subpopulations (LYVE1^hi^ also called TLF^+^ and MHCII^hi^-(CCR2^+^ or CCR2^−^)) consistently coexist across several tissues in specific sub-tissular niches ^53,54^. Along these lines, we found that the CD206^+^CD163^+^ coMac population specifically expressed *Lyve1*, *Timd4* and *Folr2* (**Figure S12**), similar to the Lyve1^hi^/TLF^+^ subpopulation ^53,54^. Meanwhile, CD206^+^CD163^−^ coMac were devoid of the previous three genes and were highly enriched in MHCII-related genes, including a subset expressing *Ccr2* (MHCII^hi^ CCR2^+^ subpopulation), suggesting that the cochlea contains three macrophage subpopulations that are conserved across several tissues (**Figure S12**).

This study further elucidates the identity of cochlea-resident macrophages by establishing their embryonic ontogeny and self-renewal capacity. Taking advantage of embryonic fate-mapping via *in utero* labeling of transgenic reporter mice to trace cells derived from the yolk sac, we demonstrated that cochlear macrophages partly originate from yolk sac precurors during development; this finding is in line with observations by Kishimoto and colleagues ^8^. In addition, our new finding of the low level of YFP positivity in cochlear macrophages compared to brain microglia suggests the existence of additional contributions, from fetal monocytes, bone marrow– derived monocytes after birth, or both ^13,22^. All compartments had a significant contribution from the primitive hematopoiesis of the yolk sac, except the spiral limbus. The functional significance of this difference in origin of macrophages within the cochlea has yet to be fully elicited and thus requires further study.

Fate-mapping analysis at adult stages showed a significant contribution from blood-derived monocytes over time restricted to the spiral lamina and spiral limbus compartments that was more pronounced in the basal and middle turns and absent in the apical turn. These findings were independently confirmed via a parabiosis model, with only GFP^+^ cells infiltrating from the blood circulation observed in the spiral lamina and the spiral limbus. Of note, we found that the pool of YFP^−^ monocytes/macrophages originating from the blood did not increase over an extended period of time (between 8 weeks and 26 weeks post tamoxifen injection) in the spiral ligament and the spiral limbus. This could suggest that blood-derived monocytes cannot compete with resident macrophages once replacement reaches a certain threshold, as previously suggested, or that only a subset of macrophages is being replaced ^55^. Our data suggest that both CD206^+^CD163^−^ coMac and CD206^+^CD163^+^ coMac subpopulations were partially renewed overtime by blood monocytes, while coMGL represented long-lived tissue-resident macrophages within the cochlea.

Here, we noted that even at 26 weeks p.i. most of the CD206^+^CD163^−^ and CD206^+^CD163^+^ coMac subpopulations were still resident YFP^+^ macrophages, thereby confirming that only a subset of cochlear macrophages is replaced by blood monocytes over time. Therefore, it is plausible that CD206^+^CD163^−^ and CD206^+^CD163^+^ coMac populations are dynamic populations, with only a limited number of available niches for blood derived monocytes. Previous studies using irradiation complemented with labeled bone marrow transfer, known to promote an artificial infiltration of blood-derived monocytes; have suggested more significant contribution of blood monocytes to the cochlea macrophage pool over time ^9,10^. Our results contrast with these previous claims and show a limited contribution of blood-derived monocytes only in specific compartments in steady state conditions. It is important to note that emerging findings in the CNS highlighted the skull bone marrow as a source of leukocytes for the CNS parenchyma in neuroinflammatory conditions ^56^. While it is unlikely such event would occur during homeostasis in the cochlea, our technical paradigm did not allow us to assess such contribution. Therefore, it remains to be determined if the bone marrow contained in the encapsulating temporal bone could supply the cochlea with monocytes that could further differentiate into macrophages during homeostasis or in pathological conditions.

Our analysis of macrophages in aged mice demonstrated that cochlear macrophages change their morphology to become larger and less ramified. Previous studies have demonstrated similar findings by investigating overall macrophage numbers and morphology without deciphering potential differences across our newly identified cochlear macrophage subpopulations ^37,57^. To gain more insights into the changes of macrophages in the aging cochlea, we examined 7-, 36-, and 70-weeks-old C57BL6/J mice. Our results demonstrate that the densities of IBA1^+^ cells increase significantly in the spiral lamina throughout the lifespan. However, we did not observe significant changes in the number of macrophages in other compartments. One possible explanation for the density increase of macrophages in the spiral lamina is that the density increase of macrophages is due to an expansion in either coMGL or sciatic nerve-like macrophages subpopulations or both. To address this question, we examined different time points of age and found that the proportions of sciatic nerve macrophage-like CD206^+^ macrophages among IBA1^+^ cells increases until 70 weeks only in the spiral lamina, while TMEM119-expressing microglia-like cell density did not change significantly. Our data suggest the contribution of each macrophage subset to the increased macrophage number in the aging cochlea differ based on the spatial localization within a tissue.

Our data suggest that cochlear tissue-resident macrophages are one of the most diverse macrophage populations in the body, with distinct molecular signatures, alluding to potentially distinct tissue-specific functions in homeostasis and disease (**Figure S13**). Their complex roles in immune regulation put them in the limelight as specific cellular targets for the therapeutic intervention of hearing loss.

## Acknowledgements

We thank the members of the Ajami, Wieghofer and Cheng labs for helpful comments on the manuscript. We thank the Flow Cytometry core, Advanced Light Microscopy core, Gene Profiling/RNA and DNA Service Shared Resources, and Massively Parallel Sequencing Shared Resources at OHSU for their support. The authors would like to thank Angela Ehrlich, Heidrun Kuhrt and Constanze Hobusch for their excellent technical assistance. Funding: OHSU institutional support (BA), R21DC021275-01 (BA), Collins Medical Trust (AChiot), Chi-Li Pao Foundation (AChing), German Academic Scholarship Foundation (MJF, DDR), NIDCD/NIH RO1DC021110 (AGC), K24DC020986 (AGC), RO1DC016919 (AGC), Medical Faculty of the University of Augsburg (PW)

## Authors contributions

Conceptualization: AC, MJF, DB, PW, BA. Investigation: AC, MJF, DB, KHR, AB. Visualization. AC, MJF, DB, KHR, AC, PW. Consultation for computational analysis: KS. Resources and discussion DDR, TM, PJA, MP and AGC. Writing – original draft: AC, MJF, DB, PW and BA. Writing – review and editing: All authors. Supervision: PW and BA

## Materials and Methods

### Mice

C57BL/6J, *Cx3cr1^CreERT^*^2^, *Rosa26-YFP*, *Tmem119*-GFP and *Ubc*-GFP mice were all purchased from The Jackson Laboratory and further bred in house. Breeding was performed on a C57BL/6J background and in pathogen-free conditions. *Cx3cr1^CreERT^*^2^ mice were crossed to *Rosa26-YFP*, and *Cx3cr1^CreERT^*^2^ (Cre/+):*Rosa26-YFP* (fl/fl) were identified by PCR screening. All animal experiments were approved by and performed in compliance with the National Institute of Health guidelines of the Institutional Animal Care and Use Committee at Oregon Health and Science University. All animals were housed under a 12-hour light cycle with water and food *ad libitum*.

### Tamoxifen injection

To induce the recombination and expression of YFP in *Cx3cr1^CreERT^*^2^*:Rosa26-YFP* mice, 6-week-old mice were injected subcutaneously with 4 mg of tamoxifen (T5648-1G, Sigma-Aldrich) dissolved in corn oil (C8267, Sigma-Aldrich) at 20 mg/ml. Three injections were performed, one every two days. To confirm recombination, blood was collected from the retro-orbital sinus one week after the last injection, and YFP expression was assessed by flow cytometry (see below). At 2-, 8-or 26-weeks post-injections, mice were deeply anesthetized with ketamine (West-Ward Pharmaceuticals) and transcardially perfused with 0.1 M ice-cold phosphate buffered saline (PBS). Blood and one cerebral hemisphere per mouse were collected to assess YFP recombination efficiency by flow cytometry in blood cells and microglia. The bone-encapsulated cochlea in the skull– and the remaining cerebral hemisphere were collected and post-fixed in 4% paraformaldehyde for 1h at room temperature (RT) for histological procedures.

### Pulse labeling

For YFP expression in yolk sac macrophages, *Cx3cr1CreERT2*:*Rosa26-YFP* pregnant females carrying embryos at E9.5, underwent i.p. injections with 75 mg/kg 4-OH-TAM (H7904, Merck) and 37.5 mg/kg Progesterone (P0130, Merck) dissolved in corn oil. At E19.5, pups were delivered by cesarean and given to foster mother mice.

### Parabiosis

6– to 7-week-old pairs of weight-matched C57BL/6J and *Ubc*-GFP mice were surgically joined as previously described (Ajami 2007). Prior to surgery, mice were housed together for two weeks and provided with soft food. To confirm blood sharing, blood was collected from the tail vein, and GFP expression was assessed by flow cytometry two weeks post-surgery. At 4– or 28-weeks post-surgery (i.e. 2 weeks and 26 weeks after blood sharing establishment), mice were deeply anesthetized with ketamine (West-Ward Pharmaceuticals) and transcardially perfused with 0.1 M ice-cold PBS. Blood, spleen and one cerebral hemisphere per mouse were collected to assess GFP^+^ cell percentage by flow cytometry. The bone-encapsulated cochlea and remaining cerebral hemisphere were collected and post-fixed in 4% paraformaldehyde for 1 h at RT for histological procedures.

### Cell preparation for flow cytometry analysis

#### For isolation of microglial cells

Brains were dissected and cut in half along the longitudinal fissure. One cerebral hemisphere per animal was homogenized with a dounce homogenizer in 0.1 M HBSS, filtered through a 70 µm cell strainer (Falcon) and centrifuged (750g, 7 min, 4°C) (Thermo Scientific). To remove the myelin, cells were resuspended in 5 ml of 0.1 M HBSS (Gibco) + 2 ml of RPMI (Gibco) + 3 ml of isotonic Percoll (ISP; 90% Percoll (GE Healthcare) + 10% 1 M HBSS (Gibco)). A 2-ml layer of 70% Percoll (70% ISP, 30% 0.1 M HBSS) was added at the bottom of the tube using spinal needles (BD Biosciences). Cells were centrifuged at 500 *g* for 15 min at RT without break. Following centrifugation, myelin was removed from the top layer, and cells were recovered from the interface between the 30% and 70% Percoll layers. Cells were washed twice using 0.1 M PBS (Gibco) supplemented with 2% fetal bovine serum (FBS; Atlas Biologicals) followed by antibody staining.

#### For isolation of splenocytes and blood cells

Spleens were dissected and homogenized with a douncer in 0.1 M HBSS and centrifuged at 750g for 7 min at 4°C. Blood was collected in a heparin tube (BD Vacutainer) either from the retro-orbital sinus or from cardiac puncture if the mouse was euthanized. Approximately 50 µl of blood was transferred to a 15-ml conical tube. Erythrocytes from spleen and blood samples were lysed in 2 ml of 10X red cell blood (RBC) lysis buffer diluted 1:10 in water (eBioscience) for 7 min at RT, according to the manufacturer’s instructions. Spleen and blood samples were then washed two times in 0.1 M PBS supplemented with 2% FBS, followed by staining.

### Flow cytometry

#### For the evaluation of the recombination efficiency in Cx3cr1^CreERT^^2^: Rosa26-YFP mice (1-, 2-, 8– and 26-weeks post injections)

White blood cell and microglial cell Fc-receptors were blocked using a purified anti-CD16/CD32 antibody (1:200, clone 93, BioLegend) for 20 min at 4°C. White blood cells were stained for 30 minutes at 4°C using a viability dye (1:2000, Ghost Dye Red 780, TONBO) in addition to the following antibodies: CD11b-BV421 (1:200, clone M1/70, BioLegend), CD115-BV605 (1:200, BioLegend, clone AFS98), CD45-BUV395 (1:200, clone 30-F11, BD Horizon) and Ly6C-PE (1:16000, clone HK1.4, BioLegend). Brain microglial cells were stained for 30 minutes at 4°C using a viability dye (1:2000, Ghost Dye Red 780, TONBO) in addition to the following antibodies: CD11b-BV421 (1:200, clone M1/70, BioLegend), CD45-BUV395 (1:200, clone 30-F11, BD Horizon).

#### For the evaluation of blood sharing in parabiotic mice (4– and 28-weeks post-surgery)

White blood cell, splenocyte and microglial cell Fc-receptors were blocked using a purified anti-CD16/CD32 antibody (1:200, clone 93, BioLegend) for 20 minutes at 4°C. White blood cells and splenocytes were then stained for 30 minutes at 4°C, using a viability dye (1:2000, Ghost Dye Red 780, TONBO) in addition to the following antibodies: CD11b-BV421 (1:200, clone M1/70, BioLegend), CD45-BUV395 (1:200, clone 30-F11, BD Horizon), CD19-BUV737 (1:1600, clone 1D3, BD Horizon), CD3ε-APC (1:400, clone 145-2C11, BioLegend) and Ly6G-PE (1:200, clone 1A8, BioLegend). Brain microglial cells were stained for 30 minutes at 4°C using a viability dye (1:2000, Ghost Dye Red 780, TONBO) in addition to the following antibodies: CD11b-BV421 (1:200, clone M1/70, BioLegend), CD45-BUV395 (1:200, clone 30-F11, BD Horizon). Cells were acquired on a BD FACSymphony Cell Analyzer (BD Biosciences), and a multiparameter analysis was performed using FlowJo Software (BD Biosciences).

### Histology and microscopy

After transcardial perfusion with PBS, skull bases were fixed in 4% PFA for 1 h at RT. Temporal bones including the cochlea were removed as described in Supplementary Figure2 dissection and decalcified in 10% EDTA for 72 h at RT. Cochlea were dissected as shown in supplementary Figure 3 dissection for whole mounts or dehydrated in 10%, 20% and 30% sucrose (step-wise raise in percentage after equilibration of the tissue) and embedded in Tissue-Tek O.C.T.TM Compound for cryosections.

#### Whole mounts

whole mounts were blocked with PBS containing 1% bovine serum albumin, 0.3% Triton X-100 for permeabilization and 2% normal donkey serum (NDS; if secondary antibody species was donkey) or 2% normal goat serum (NGS; if secondary antibody species was goat) for 24 h at 4°C. Primary antibodies were added overnight (**Table S1**) at 4°C in a blocking solution. After washing 7x for 45 min at RT with a washing solution containing 0.2% NDS or NGS, secondary antibodies were added overnight at 4°C. Secondary antibodies were washed 7x for 45 min at RT with washing solution (see antibody table for references and dilutions), and nuclei were counterstained with 4’,6-Diamidino-2-phenylindole (DAPI). In the case of staining with CD206-Alexa Fluor^®^-647 antibody was added over an additional night at 4°C in blocking solution and then washed and counterstained as previously described. Whole mount confocal pictures were taken with a Fluoview FV1000 (Olympus) using a 20x 0.75 NA U Plan S Apo, or a 40x 0.95 NA U Plan S Apo, or a Zeiss LSM 980 using a 10x 0.45 M27 Plan Apo.

#### Cryosections

12 µm cryosections were blocked with PBS containing 2% bovine serum albumin and 0.1% Triton X-100 for permeabilization and 2% normal donkey serum (NDS; if secondary antibody species was donkey) or 2% normal goat serum (NGS; if secondary antibody species was goat) for 2 h at RT. Primary antibodies were added overnight at 4°C in a blocking solution (**Table S1**). After washing 3x for 20 min at RT with a washing solution containing 0.2% NDS or NGS, secondary antibodies were added for 2h at RT in the blocking solution. After washing 3x for 20 min at RT with washing solution, nuclei were counterstained with 4’,6-Diamidino-2-phenylindole (DAPI). Cryosection images were taken using a conventional fluorescence microscope (Olympus BX-40) with a color camera (Olympus XM-10).

#### Immunofluorescence Analysis

The lack of bloodstream replenishment of microglial cells during homeostasis has been demonstrated, hence YFP expression levels are expected to be stable in long-lived tissue macrophages. To normalize variability induced by the tamoxifen injections in each mouse, the percentage of YFP-expressing macrophages in the cochlea was normalized with the individual and corresponding percentage of brain YFP+ microglia from the same animal, determined by flow cytometry (see above).

### Tissue preparation for sorting

Each mouse was anesthetized using ketamine (West-Ward Pharmaceuticals) and then transcardially perfused with 20 mL of ice-cold 0.1 M PBS prior to tissue harvest. Three mice were pooled for each sample of brain, spleen, lung, peritoneal macrophages and liver. Brain and spleen samples were prepared as described in the flow cytometry section (see above). Lungs were collected, homogenized using a douncer in 0.1 M HBSS, filtered through a 70 µm cell strainer and washed two times in 0.1 M PBS + 2% FBS prior to staining. For peritoneal macrophages, mice were injected twice with 5 ml of ice-cold 0.1 M HBSS in the peritoneal cavity. Peritoneal macrophages were harvested by carefully recollecting the fluid from the peritoneum. Cells were washed two times with 0.1 M PBS supplemented with 2% FBS before staining. For the liver, one lobe of liver per mouse was collected. Lobes were homogenized using a douncer and filtered through a 70 µm cell strainer. Liver tissues were then enzymatically digested in a 0.1 M HBSS solution containing 2 mg/ml Dispase II (Sigma Aldrich), 4 mg/ml Collagenase IV (Worthington) and 15 mM HEPES media (Gibco) on a shaker for 45 minutes at 37°C. Following incubations, cells were washed one time in 0.1 M HBSS, resuspended in 15 ml of 35% Percoll and centrifuged for 15 min at 2200 rpm with no break at RT. Cells were collected from the bottom of the tube and washed two times in 0.1 M PBS supplemented with 2% FBS prior to staining.

Seven mice were pooled for retina, sciatic nerve, and cochlea samples. For the retina, eyes were dissected under a dissecting microscope to harvest retinas. Retinas were dissociated by resuspending the tissue using P1000 and P200 tips. Post-dissociation, retinas were filtered through a 70 µm cell strainer and washed two times in 0.1 M PBS supplemented with 2% FBS before staining. For cochlea, cochlea were detached from the temporal bone, and cochlea were carefully dissected out of the bone shell under a dissecting microscope. The pool of cochlea then underwent mechanical dissociation by cutting prior to resuspension in enzymatic digestion buffer (0.1 M HBSS solution containing 2 mg/ml Dispase II (Sigma Aldrich), 4 mg/ml Collagenase IV (Worthington) and 15 mM HEPES media (Gibco)) in a 2 mL Eppendorf tube. For sciatic nerves, left and right nerves from the gastrocnemius muscle to the vertebral column were collected. Two sciatic nerves per 2 mL Eppendorf tube were mechanically dissociated and then resuspended in enzymatic digestion buffer (0.1 M HBSS solution containing 2 mg/ml Dispase II (Sigma Aldrich), 4 mg/ml Collagenase IV (Worthington) and 15 mM HEPES media (Gibco)) in a 2 mL Eppendorf tube. Cochlea and sciatic nerve tubes were then incubated on a shaker at 37°C for 45 min. Following enzymatic digestion, cells were resuspended once again using P200 tips, filtered using a 70 µm cell strainer and washed two times with 0.1 M PBS + 2% FBS prior to staining.

### Cell Sorting

Each tissue replicate is an independent experiment for a total of 3-4 experiments per tissue (Supplementary Figure 1). Cell Fc receptors were blocked using a purified anti-CD16/CD32 antibody (1:200, clone 93, BioLegend) for 20 min at 4°C. Cells were then stained for 30 min at 4°C, using a viability dye (1:200, Ghost Dye Red 780, TONBO) in addition to specific panels of antibodies for each tissue. Brain: CD11b-BV421 (1:200, clone M1/70, BioLegend), CD45-BUV395 (1:200, clone 30-F11, BD Horizon), F4/80-APC (1:200, clone BM8, BioLegend). Spleen: CD11b-BV421 (1:200, clone M1/70, BioLegend), CD45-BUV395 (1:400, clone 30-F11, BD Horizon), Ly6G-PE (1:200, clone 1A8, BioLegend), Ly6C-PE (1:16000, clone HK1.4, BioLegend), CD19-BUV737 (1:200, clone 1D3, BD Horizon), F4/80-APC (1:200, clone BM8, BioLegend). Lung: CD11b-BV421 (1:200, clone M1/70, BioLegend), CD45-BUV395 (1:400, clone 30-F11, BD Horizon), Ly6G-PE (1:200, clone 1A8, BioLegend), Ly6C-PE (1:16000, clone HK1.4, BioLegend), CD11c-FITC (1:1600, clone N418, BioLegend), CD24-APC (1:400, clone M1/69, BioLegend), CD64-PE/Cy7 (1:200, clone X54-5/7.1, BioLegend). Peritoneal macrophages: CD11b-BV421 (1:200, clone M1/70, BioLegend), CD45-BUV395 (1:200, clone 30-F11, BD Horizon), CD19-BUV737 (1:200, clone 1D3, BD Horizon), F4/80-APC (1:200, clone BM8, BioLegend). Liver: CD11b-BV421 (1:200, clone M1/70, BioLegend), CD45-BUV395 (1:400, clone 30-F11, BD Horizon), F4/80-APC (1:200, clone BM8, BioLegend), Ly6G-PE (1:200, clone 1A8, BioLegend), CD19-BUV737 (1:200, clone 1D3, BD Horizon). Retina and Sciatic nerve: CD11b-BV421 (1:200, clone M1/70, BioLegend), CD45-BUV395 (1:200, clone 30-F11, BD Horizon), F4/80-APC (1:200, clone BM8, BioLegend), Ly6G-PE (1:200, clone 1A8, BioLegend), Ly6C-PE (1:16000, clone HK1.4, BioLegend), CD64-PE/Cy7 (1:200, clone X54-5/7.1, BioLegend). Cochlea: CD11b-BV421 (1:200, clone M1/70, BioLegend), CD45-BUV395 (1:200, clone 30-F11, BD Horizon), F4/80-APC (1:200, clone BM8, BioLegend), Ly6G-PE (1:200, clone 1A8, BioLegend), Ly6C-PE (1:16000, clone HK1.4, BioLegend). Cells were sorted with a FACSAria Fusion Cell Sorter (BD Biosciences) or a FACSymphony S6 Cell Sorter (BD Biosciences) using a 130 µm nozzle. For each sample, between 3500 and 6000 live macrophages were sorted into 0.1 M PBS containing 0.04% BSA. Cells were maintained at 4°C for the entire experiment.

### Single-cell sequencing

Cells were processed according to the Chromium Next GEM Single-cell 3’ v3.1 protocol from 10X Genomics (CG000315 Rev B). Immediately after sorting, cells were centrifuged at 500 *g* for 10 min and resuspended in 43.1 µl of 0.1M PBS containing 0.04% BSA. In parallel, reverse transcription master mix was prepared according to manufacturer instructions, and 31.9 µl of master mix was added to the cells. The cell suspension was loaded into the Chromium (10X Genomics) to obtain Gel beads in EMulsion (GEMs), each containing a single-cell, a gel bead with barcoded oligonucleotides and reverse transcription reagents. Following GEM generation, reverse transcription was performed by incubating the GEMs in a thermal cycler at 53°C for 45 min and 85°C for 5 min and then the GEMS were stored at –20°C for one week maximum until cDNA amplification. GEM-RT product clean-up was performed using Dynabeads MyOne Silane beads (ThermoFisher) to isolate nucleic acid. cDNA was amplified according to the following protocol: 98°C for 3 min; 12 or 13 cycles of 98°C for 15 s, 63°C for 20 seconds and 72°C for 1 minutes; 72°C for 1 minutes and 4°C on hold. Amplified cDNA was cleaned up using SPRIselect reagents (Beckman Coulter). cDNA was stored for a maximum of 4 weeks at –20°C before making the libraries and concentration was assessed using a Bioanalyzer High Sensitivity DNA Kit on a 2100 Bioanalyzer (Agilent). To generate libraries, 10 µL cDNA (1/4 of the total cDNA obtained from the amplification/clean-up steps) was used. A pre-cooled block at 4°C was required for the DNA fragmentation, end-repair and A-tailing, and samples were incubated at 32°C for 5 min and 65°C for 30 min. A double-sided size clean-up of the fragments was performed using SPRIselect beads. Samples were then incubated at 20°C for 15 min for the adaptor ligation and then cleaned up with SPRIselect beads. Sample Index PCR was performed according to the following protocol: 98°C for 45 s; 13-16 cycles (depending on cDNA input) of 98°C for 20 s, 54°C for 30 s, 72°C for 20 s and 72°C for one min prior to holding at 4°C. Each sample was indexed to a unique set of two indices in order to run all the samples in a multiplexed sequencing run. A final double-sided size selection was performed using SPRIselect beads. Library traces were obtained using a Bioanalyzer High Sensitivity DNA Kit on a 2100 Bioanalyzer (Agilent). Libraries were stored at –20 for long-term storage until sequencing. RNA sequencing was performed by the Massive Parallel Sequencing Shared Resource (MPSSR) at OHSU. Paired-end sequencing was performed on a NovaSeq 6000 (Illumina) at the depth of 50,000 reads per cell. The demultiplexing of the data into FASTQ files was performed using Cell Ranger mkfastq pipeline (10X Genomics). The Cell Ranger count pipeline was used to perform alignment of the data on the mouse reference genome (mm10), filtering and barcode and UMI counting.

### Single-cell RNA-seq analysis

#### Data Preprocessing/QC

The Seurat version 4.1.1 R package was used to process and visualize single-cell RNA data. Each 10x library was loaded using Seurat’s Read10X function, after which they were individually clustered and processed using the libraries DoubletFinder version 2.0.3 and SoupX version 1.6.2 for the removal of doublets and ambient RNA, respectively ^58,59^. Per-sample selection and removal of outlier cells was done manually based on 3 metrics of cell quality: proportion of reads mapping to mitochondrial genes, number of unique genes detected, and number of UMIs. An initial cell clustering was performed according to the workflow proposed in Seurat’s guided clustering tutorial in order to identify faulty populations (such as Ptprc negative cells) and batch effects present in both Retina and Spleen resident macrophage populations, which were removed from further analysis. An additional cluster of low-quality cells originating from the tissues brain and retina was also removed, likely representing low-quality microglia.

#### Cell clustering

In order to improve the biological accuracy of clustering between tissues and reduce the confounding effect of the dissociation method, a list of 218 known dissociation genes was sourced from Van Hove et al. ^23^. An additional 161 genes were found to be highly correlated (Spearman’s R > 0.3) with a gene module constructed from the aforementioned 218 known dissociation genes, collectively designated as a set of 379 ‘dissociation related’ genes. Cells were re-clustered by first finding the top 2000 most variable features among all post-qc cells using Seurat’s FindVariableFeatures function and 182 ‘dissociation-related’ genes found to be highly variable in our dataset were removed from the list of variable features. Seurat pipeline was then used to scale gene expression values using ScaleData function and the RunPCA function generated 50 principal components. Seurat’s ElbowPlot function showed that just the first 17 components explained most of the variation in the dataset, and so we chose to use them for subsequent clustering and dimensional reduction. A k = 20 nearest neighbors data object was created for the whole dataset using the Seurat function FindNeighbors, using default parameters aside from the components used. The neighbor graph object was Leiden clustered using Seurat’s FindClusters, again using the first 17 components, a resolution of 0.5, and otherwise default parameters. Finally, a Uniform Manifold Approximation and Projection (UMAP) low-dimensional embedding was generated via the RunUMAP function using default settings aside from the number of components used.

Cochlea immune cells were also clustered separately from other tissue cells, which involved first subsetting the full dataset to cochlea samples only, then finding 1500 new variable features to use for clustering. As before, gene expression values were scaled, regressing out QC metrics, cell cycle scores, and percent of counts mapping to ribosomal genes. PCA was run generating 20 PCs, the top 10 of which were used for neighbor finding, clustering, and low-dimensional projection. The resolution used in clustering was 0.3. Removal of ‘dissociation-related’ genes from the set of most variable features was found to have a miniscule effect on cochlea-only clustering, so it was not performed.

#### Cell type annotation

The reference-based cell annotation algorithm SingleR version 1.8.1 was used with the ImmGen cell type database (accessed through the package celldex 1.4.0) to make initial inferences about cell type per-cell ^60,61^. These initial annotations were improved by using Seurat’s FindMarkers function. Cross-referencing top markers found in each cluster with relevant literature led to more biologically meaningful annotations. After annotation, certain cochlea populations were chosen to be excluded from further analysis, namely 4 small clusters representing 2 populations of eosinophils, 1 population of basophils, and 1 population of B cells.

#### Gene Expression Visualization

Marker genes for each cell type were identified using Seurat’s FindMarkers and FindAllMarkers functions. Markers were visualized via ComplexHeatmap version 2.10.0, or Seurat’s VlnPlot and DotPlot functions ^62^. In heatmap visualizations, normalized gene expression values were averaged within annotated cell type groups and either rescaled by max-scaling (dividing each expression value by the maximum gene expression value) or by averaging scaled expression values within groups, then clipping averaged values between –0.5 and 0.5. The cell type correlation plot was generated using the corrplot R package version 0.92 ^63^. Pearson’s R scores were generated using a matrix of log normalized counts averaged within each cell type group. The genes selected for correlation analysis were the same used in clustering (i.e the top 2000 most variable features selected using Seurat’s FindVariableFeatures function, minus ‘dissociation genes’).

#### Downstream Analysis

Functional enrichment analysis was carried out via the clusterProfier R package version 4.2.2 ^64^. Differential expression analysis via Seurat’s FindMarkers function was used in each cell type group to identify the top 100 most exclusive markers relative to other cell type groups. Each set of markers were ordered by average log2 fold change and filtered for significance (adjusted p-value < 0.01) and average log2 fold change (avg_log2fc > 0.4). Finally, dissociation-related genes were removed from found markers prior to enrichment. Each set of filtered markers was supplied to clusterProfiler’s compareCluster function, and gene set enrichment was calculated for each via enrichGO: clusterprofiler’s implementation of an overrepresentation test. Gene sets used for enrichment were selected from GO ontology, biological processes available through the org.Mm.eg.db package version 3.14.0 ^65^ and subsetted to gene sets between 10 and 350 gene sets in length. ClusterProfiler’s implementation of GOSemSim, accessed through the simplify function, was used to simplify results using a similarity cutoff of 0.7 ^66^. Enriched gene sets for each cell type group were ordered by qvalue, and the top 30 for each group were displayed using clusterProfiler’s emapplot function. Gene sets of interest were selected from the top 20 most enriched gene sets in each group for further visualization using ComplexHeatmap. Transcription factor absolute expression was generated by simply filtering the resulting genes of Seurat’s FindAllMarkers for known mouse transcription factors, based on previously established human transcription factors ^67,68^. PCA plots were generated by averaging the first 2 PC scores within sample groups and plotting using ggplot2.

### Statistical analysis

Statistical analyses were performed using GraphPad Prism (GraphPad software, version 9.4.0 and 6.0). Data were tested for Normality using the Shapiro-Wilk normality test. If data were normally distributed, t-test with Welch’s corrections or Welch’s one-way ANOVA were performed, if not Mann-Whitney or Kruskal-Willis tests were performed. Level of significance between the three cochlear turn sections (apical-middle-basal) was tested by Repeated Measures ANOVA with geisser-Greenhouse correction in case of normal distribution, otherwise, Friedman test was performed. Data were considered significant if p-value < 0.05. See **tables S2 to S5** for a summary of the means, SEM and p-values presented in this manuscript.

