## Supplemental Information for "Single-cell, spatial, and fate-mapping analyses uncover niche dependent diversity of cochlear myeloid cells"

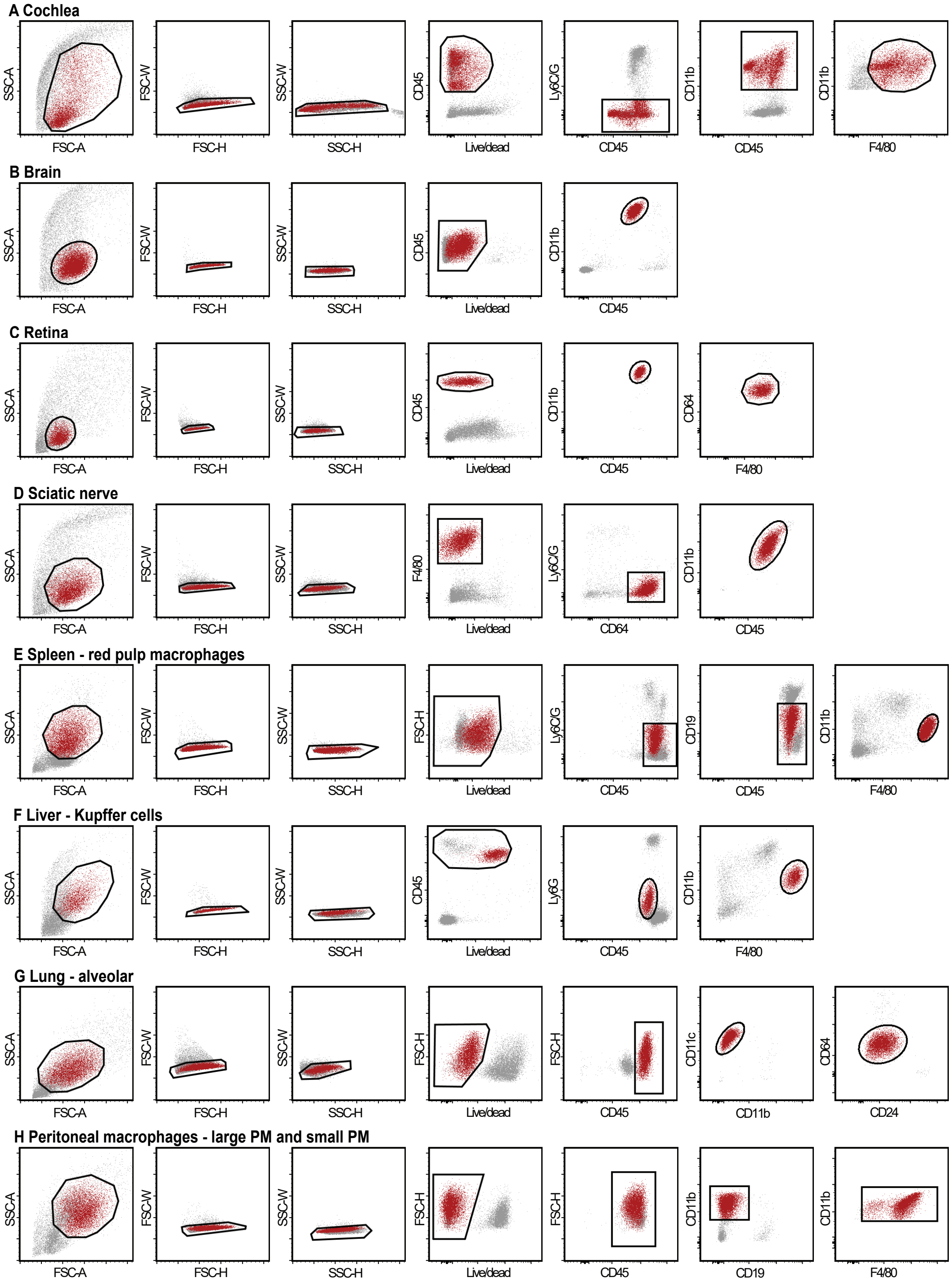


fig. S1

**Figure S1: Gating strategy for collecting macrophages from the different tissues.**

**A-H.** Overview of the gating strategy used for collecting macrophages from the cochlea (A), microglial cells from the brain (B) and the retina (C), macrophages from the sciatic nerve (D), red-pulp macrophages from the spleen (E), Kupffer cells from the liver (F), alveolar macrophages from the lung (G) and small and large peritoneal macrophages (H). Black gates show the actual gating used to collect the cells. Cells highlighted in red show the back gating of the population in previous gates.

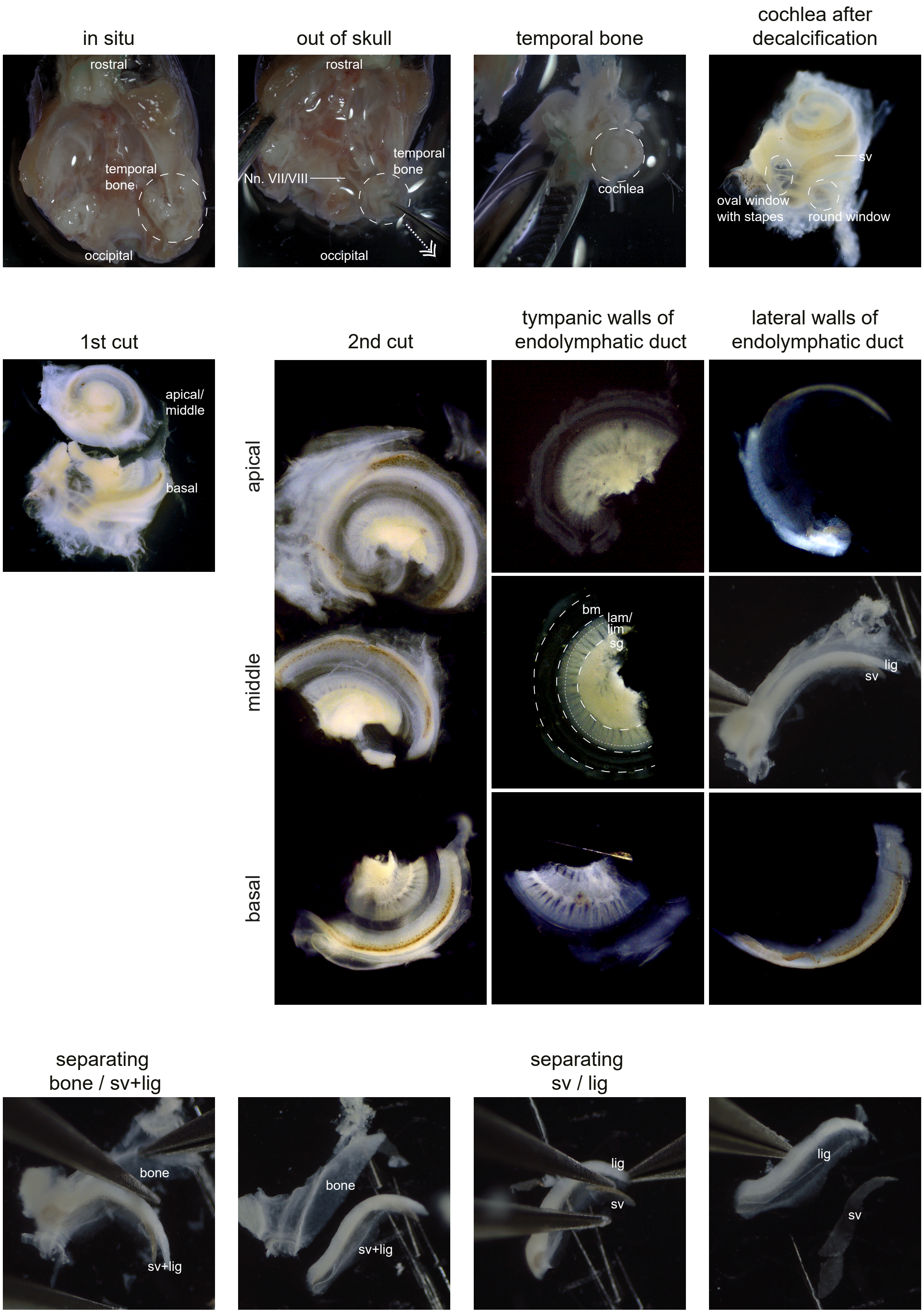
in situ
rostral
temporal bone
occipital
out of skull
rostral
Nn. VII/VIII
temporal bone
occipital
temporal bone
cochlea
cochlea after decalcification
sv
oval window with stapes
round window
1st cut
apical/ middle
basal
2nd cut
apical
middle
basal
tympanic walls of endolymphatic duct
bm
lam/ lim
sg
lateral walls of endolymphatic duct
sv
lig
separating bone / sv+lig
bone
sv+lig
bone
sv+lig
separating sv / lig
lig
sv
lig
sv

fig. S2

15

**Figure S2: Dissection and separation of cochlear compartments for immunofluorescence.** After craniotomy and removal of the brain, the temporal bone containing the cochlea is dislodged out of the skull base using a standard pattern forceps. After decalcification and cutting away excess temporal bone and otic capsule tissue using a 2.5 mm spring scissors, a cut is done along the lateral wall of the basal turn with one scissors blade in the oval window. After that, one blade is put into the opened cochlear duct and the other on the outside of the temporal bone, medial to the oval window. These first cuts separate the basal turn from the middle and apical turns. Then, cuts are placed along the lateral wall of the middle turn where it used to be connected to the basal turn with one blade inserted into the scala vestibuli. After that, one blade is put into the cut region, with the middle turn placed on top of the blade, and the other blade on the outside of the bony shell. This separates the middle turn and the apical turn from each other. After cutting between basilar membrane and lateral wall, removing Reiner's and tectorial membrane and trimming, tissue pieces of the tympanic walls of the apical, middle and basal turns consisting of basilar membrane, spiral lamina and spiral ganglion, and the lateral walls consisting of stria vascularis, spiral ligament and bony shell can be collected and separated from each other for further processing. For separating the compartments of the lateral wall, first the stria vascularis-spiral ligament compound is peeled out of the bony shell using forceps and subsequently stria vascularis and spiral ligament are pulled apart. Nn. VII/VIII: cranial nerve VII and VIII, sv: stria vascularis, bm: basilar membrane, lam: spiral lamina, lig: spiral ligament, sg: spiral ganglion.

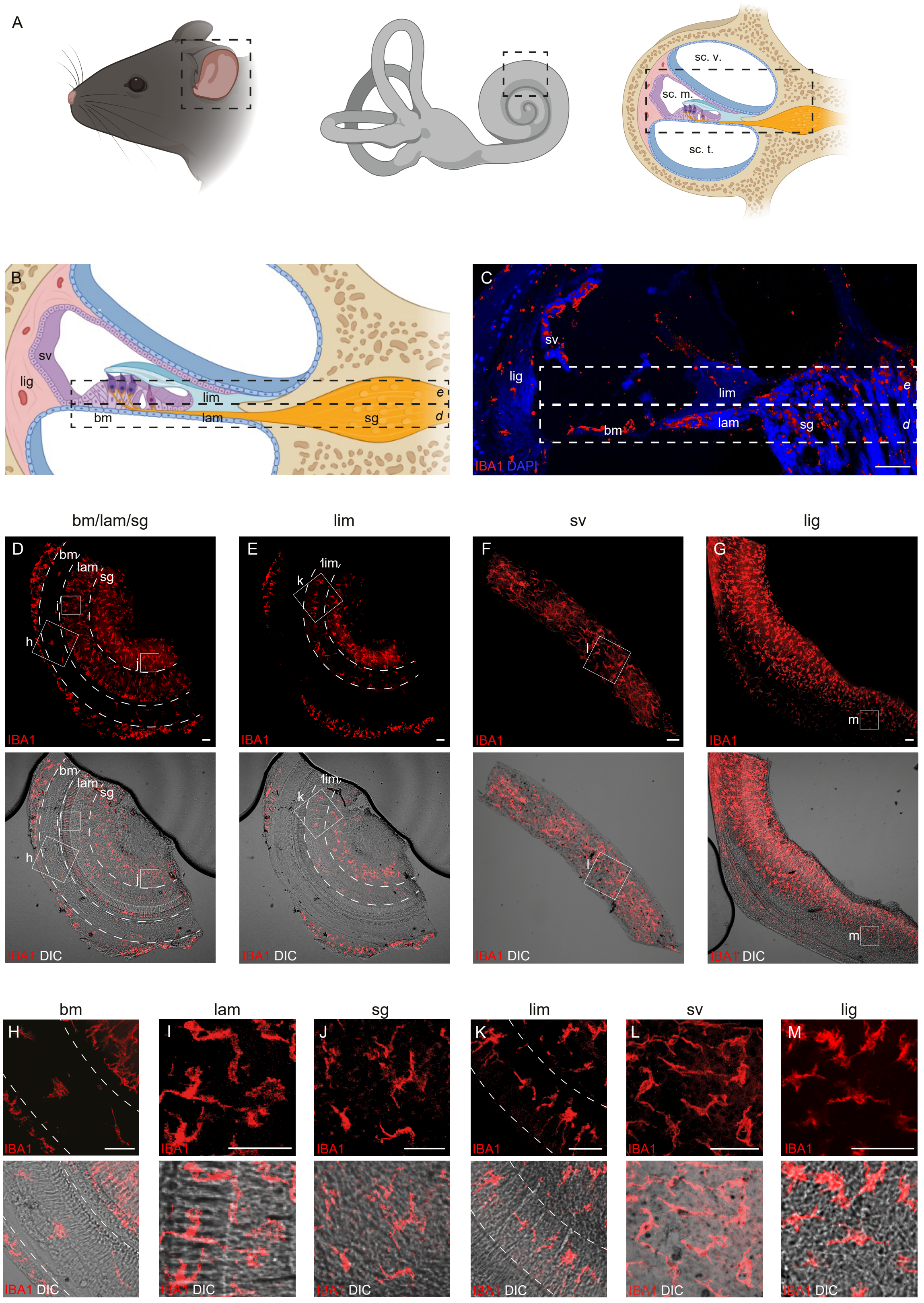

A
sc. v.
sc. m.
sc. t.
B
sv
lig
lim
lam
bm
sg
e
d
C
sv
lig
lim
lam
bm
sg
e
d
IBA1 DAPI
bm/lam/sg
lim
sv
lig
D
E
F
G
bm
lam
sg
i
h
j
k
lim
l
m
IBA1
IBA1 DIC
bm
lam
sg
lim
sv
lig
H
I
J
K
L
M


fig. S3

36

**Figure S3: Identification and localization of cochlear macrophages across compartments.**

**A.** Left & middle: anatomic localization and scheme of the inner ear consisting of the vestibular system and the cochlea. Right: Cross-sectional scheme of the cochlear ducts (from down to above: scala tympani = sc. t., scala media = sc. m. and scala vestibuli = sc. v.).

**B.** Cross-sectional scheme of the scala media with its lateral wall, consisting of stria vascularis (sv) and spiral ligament (lig), and its tympanic wall, consisting of basilar membrane (bm), spiral lamina (lam), spiral ganglion (sg) and spiral limbus (lim).

**C.** Same cross-sectional view at the scala media after cross-sectioning and fluorescence microscopy of cryo-embedded cochlea from a WT mouse. Macrophages are visualized by the pan-macrophage marker IBA1 (red). Scale bar = 100 µm.

**D-G.** Maximum intensity projections of the different cochlear compartments. Top: Macrophages are stained with IBA1 (red). Bottom: IBA1 is overlaid with differential interference contrast (DIC) to visualize the tissue by its physical light scattering properties. Scale bars = 50 µm. Maximum intensity projections of the basilar membrane (bm), spiral lamina (lam), spiral ganglion (sg) separated by a white dashed line (**D**), of the spiral limbus (lim) in between the two dashed lines (**E**), of the stria vascularis (sv, **F**) and of the spiral ligament (lig, **G**).

**H-M**. Higher magnified pictures of the basilar membrane (bm, **H**), spiral lamina (lam, **I**), the spiral ganglion (sg, **J**), the spiral limbus (lim, **K**), the stria vascularis (sv, **L**) and the spiral ligament (lig, **M**). The localization of each magnified picture is indicated by a white square in Figure S2, D to G. Each white square contains approximatively the same number of cells. Top: Macrophages are stained with IBA1 (red). Bottom: IBA1 is overlaid with differential interference contrast to visualize the tissue by its physical light scattering properties. Scale bars = 50 µm.

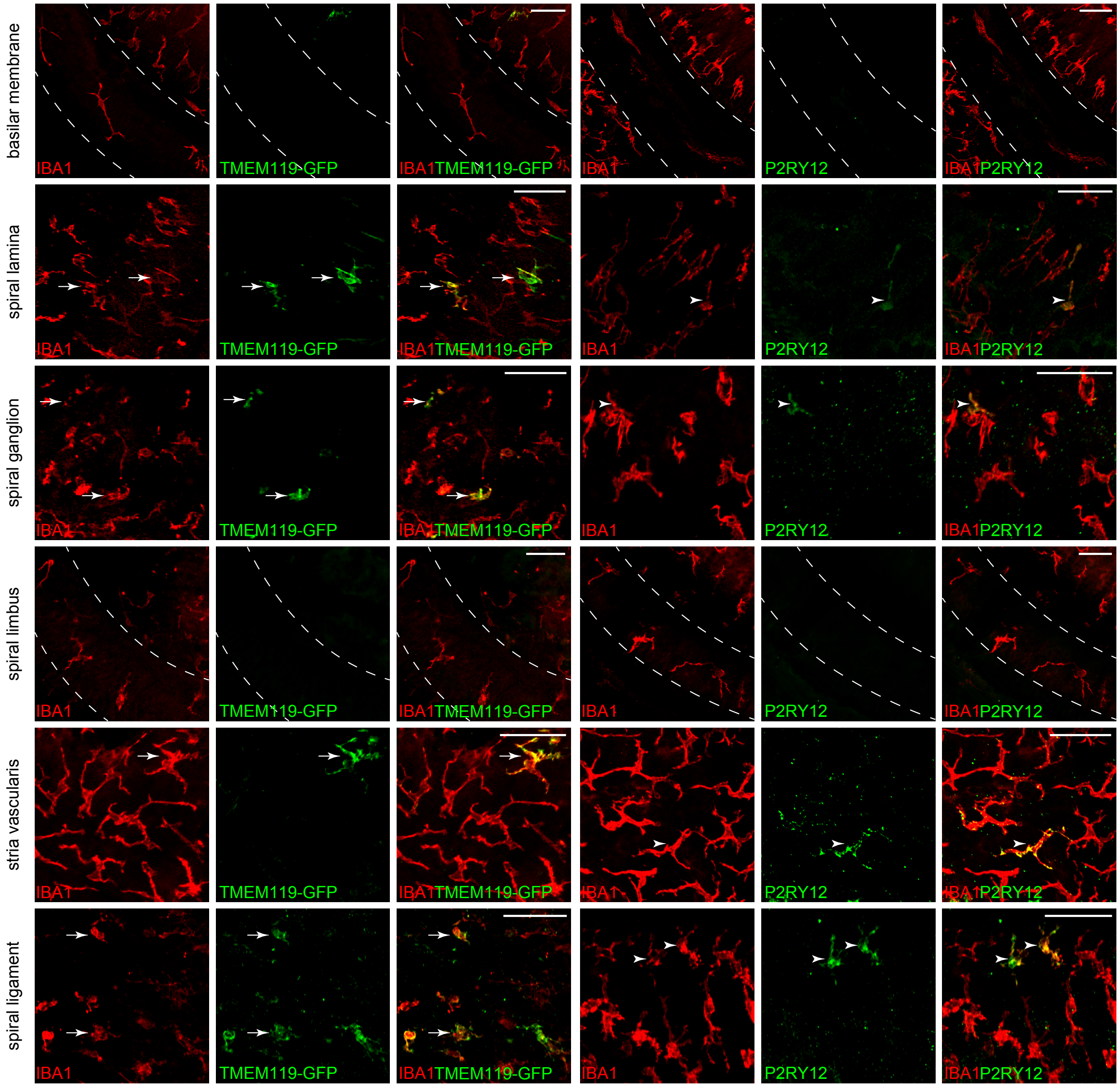


fig. S4

**Figure S4: *Tmem119*-GFP and P2RY12 marker expression across all compartments in the cochlea.**

Representative immunofluorescence images from the apical, middle, and basal segments of the basilar membrane, spiral lamina, spiral ganglion, spiral limbus, stria vascularis, and spiral ligament from whole mount cochlea of 7 weeks old *Tmem119*-GFP mice or 7 weeks old C57BL/6J mice. IBA1 is shown in red, and *Tmem119*-GFP- positive (arrows) or P2RY12-positive (arrowhead) cells in green. Representative image of n=3 animals. Scale bars represent 50 μm.

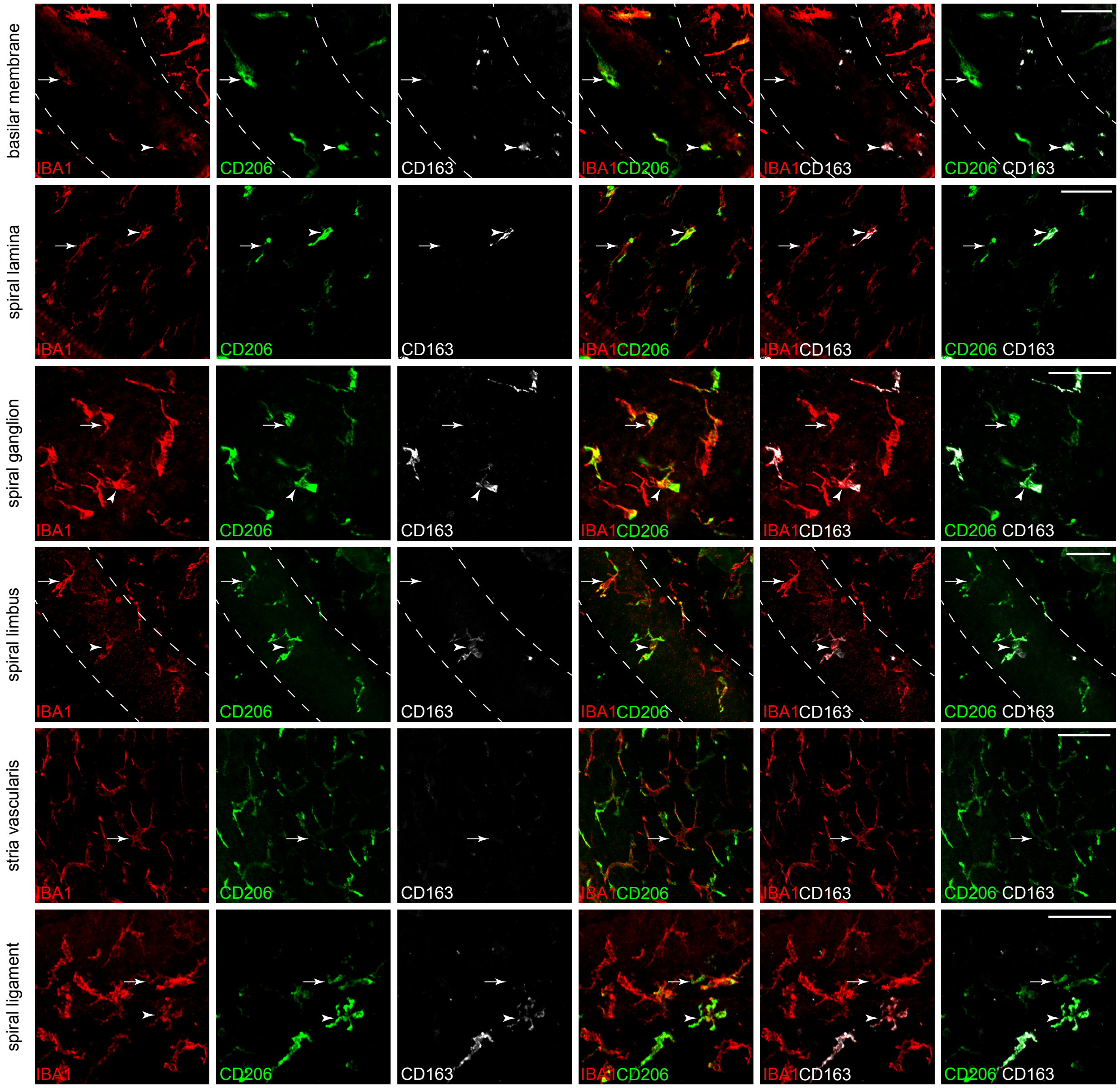


fig. S5

**Figure S5: CD206 and CD163 marker expression across all compartments in the cochlea.** Representative immunofluorescence images from the apical, middle, and basal segments of the basilar membrane, spiral lamina, spiral ganglion, spiral limbus, stria vascularis, and spiral ligament from whole mount cochlea of 7 weeks old C57BL/6J mice. Pan-macrophage marker IBA1 is shown in red, CD206 in green, and CD163 in white revealing $IBA1^+ CD206^+ CD163^-$ (arrow) and $IBA1^+ CD206^+ CD163^+$ (arrowhead) macrophages. Representative image of n=3 animals. Scale bars = 50 μm.

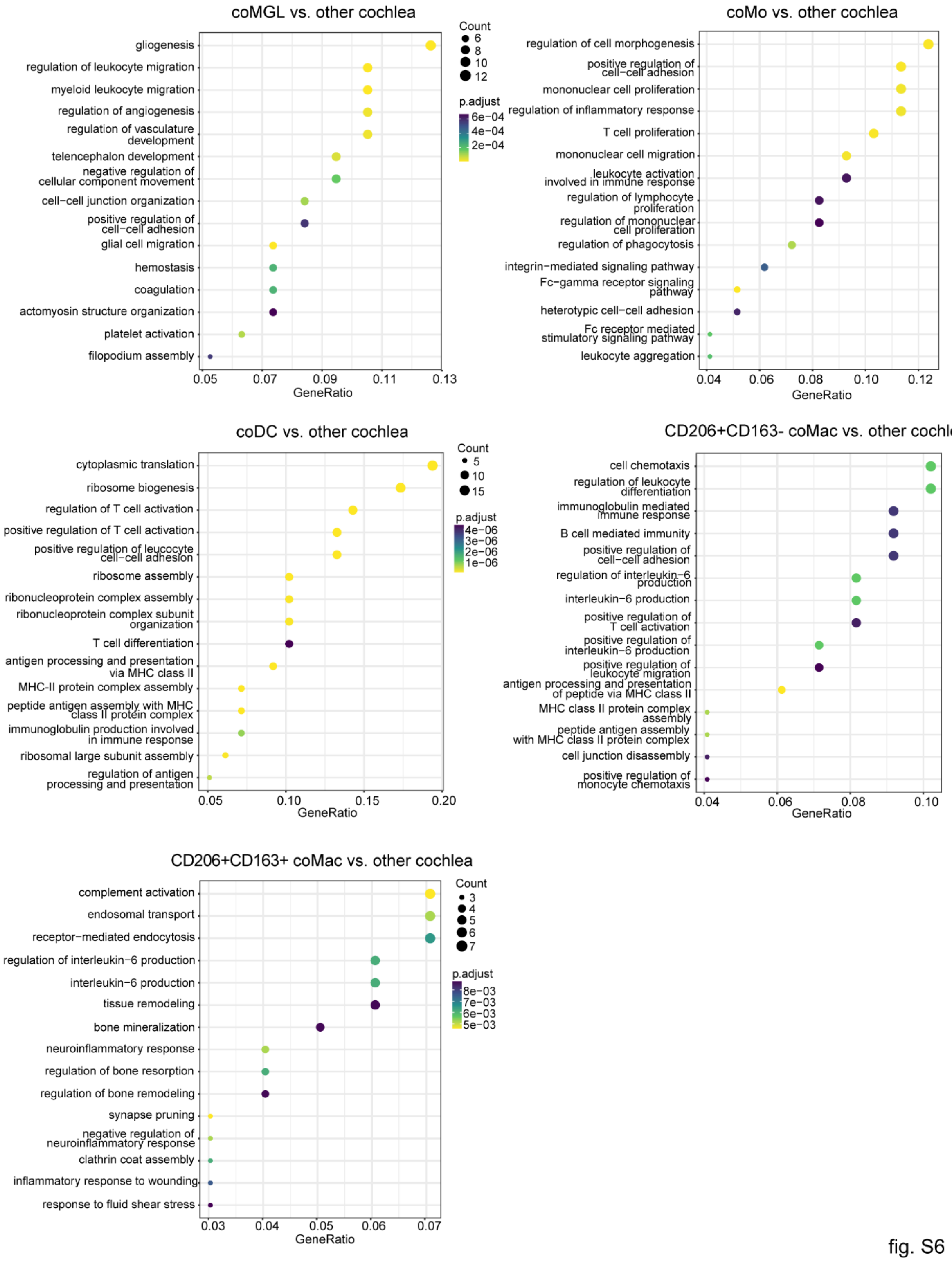


fig. S6

**Figure S6: Functional enrichment of cochlea clusters.** Dot plot showing the top 15 GO terms enriched in each cochlea cluster versus all the other cochlea clusters as a group. The size of the dot depicts gene counts for each pathway and colors display the p-value associated with each pathway.

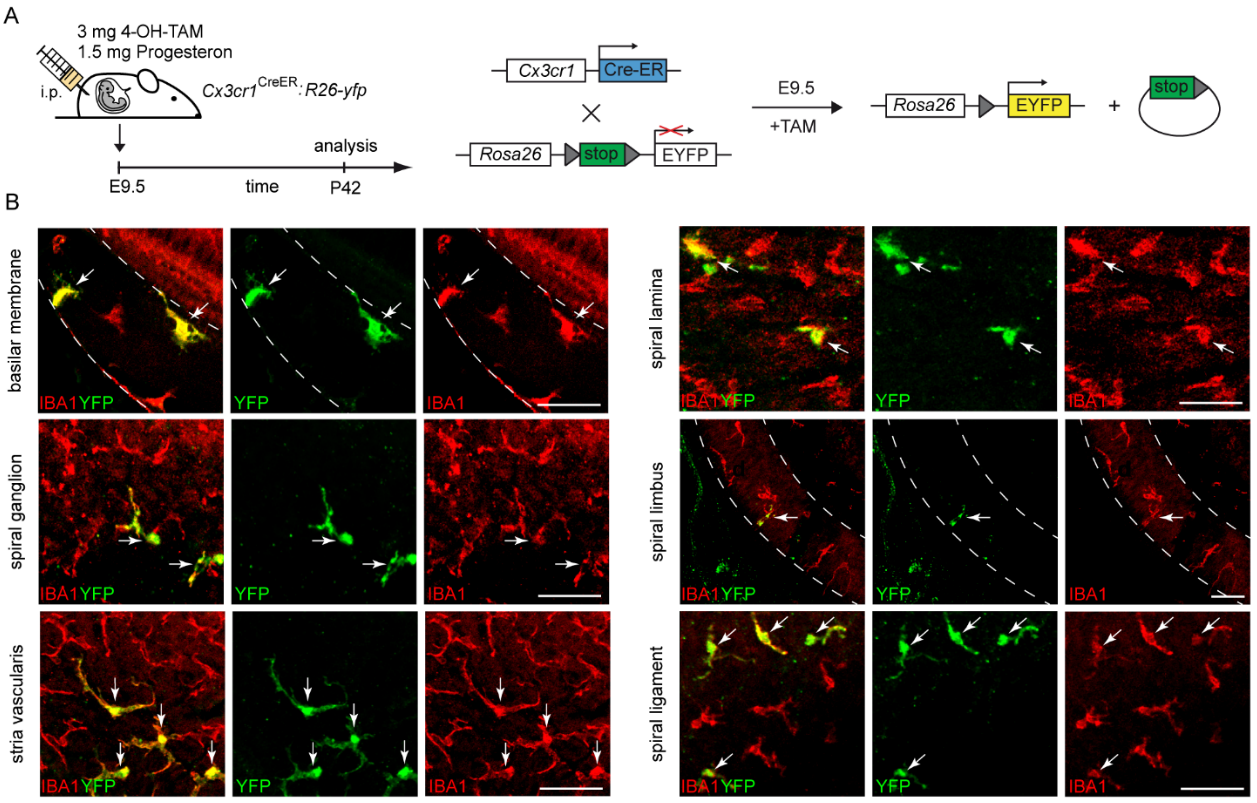


fig. S7

**Figure S7: Ontogeny of cochlear macrophages in steady-state conditions.**

**A.** Scheme of a fate-mapping experiment using *Cx3cr1*$^{CreERT2}$*:Rosa26-YFP* mice. 4-OH-Tamoxifen (4-OH-TAM) and progesterone injection were performed on embryonic day 9.5 (E9.5). Administration of 4-OH-TAM leads to intra-embryonic excision of a stop sequence flanked by loxP sites (gray triangles) in *Cx3cr1* expressing cells which causes stable and steady YFP expression under the control of the *Rosa26* promotor. Mice were evaluated on postnatal day 42 (P42).

**B.** Representative immunofluorescence images from whole mounts of the different cochlear compartments in TAM-induced *Cx3cr1*$^{CreERT2}$*:Rosa26-YFP* mice at P42. Arrows show IBA1$^{+}$ YFP$^{+}$ cells originating from the primitive hematopoiesis of the yolk sac. Scale bars = 50 µm.

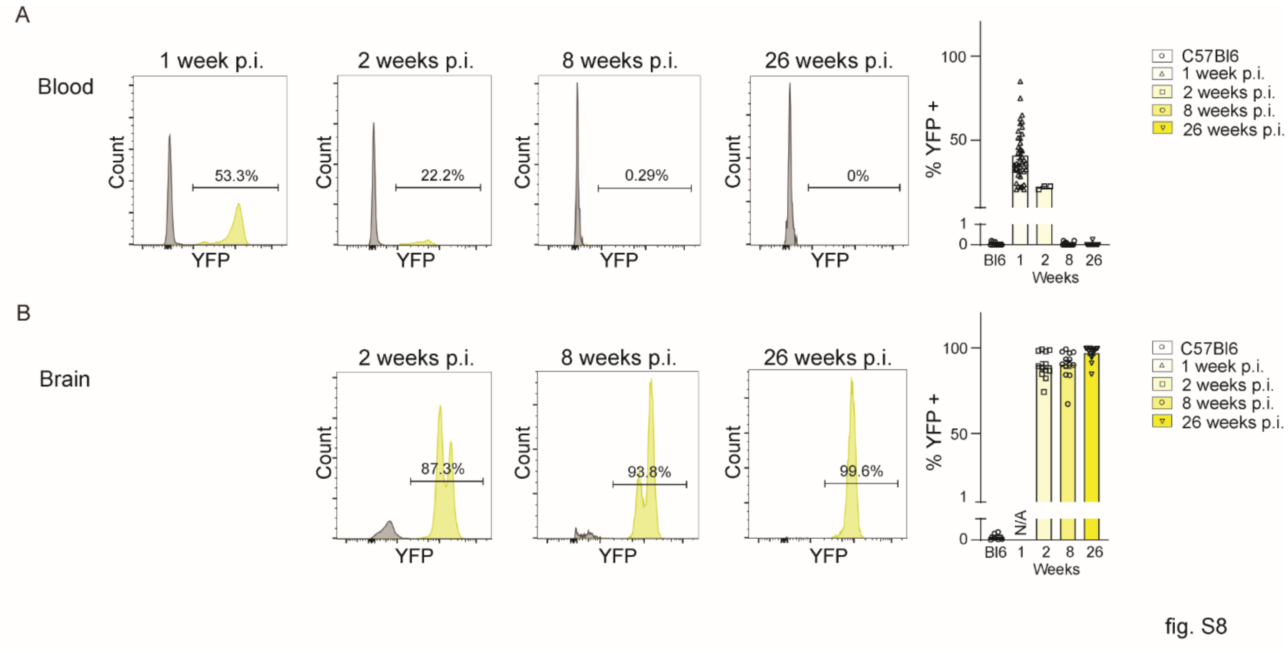


**Figure S8: Tamoxifen-induced YFP expression in *Cx3cr1*$^{\mathrm{CreERT2}}$:Rosa26-YFP mice.**

**A.** Histogram showing a representative expression of YFP by blood monocytes at different time points and quantification of the YFP expression in monocytes of control C57BL/6J (n=13) and Tamoxifen-injected *Cx3cr1*$^{CreERT2}$*:Rosa26-YFP* mice at 1-week (n=42), 2-weeks (n=4), 8 weeks (n=16) and 26 weeks (n=10) post tamoxifen injection (p.i.).

**B.** Histogram showing a representative expression of YFP by brain myeloid cells at the different time points and quantification of YFP expression by brain myeloid cells of control C57BL/6J (n=6) and Tamoxifen-injected *Cx3cr1*$^{CreERT2}$*:Rosa26-YFP* mice at 2 weeks (n=12), 8 weeks (n=13) and 26 weeks (n=14) post tamoxifen injection.

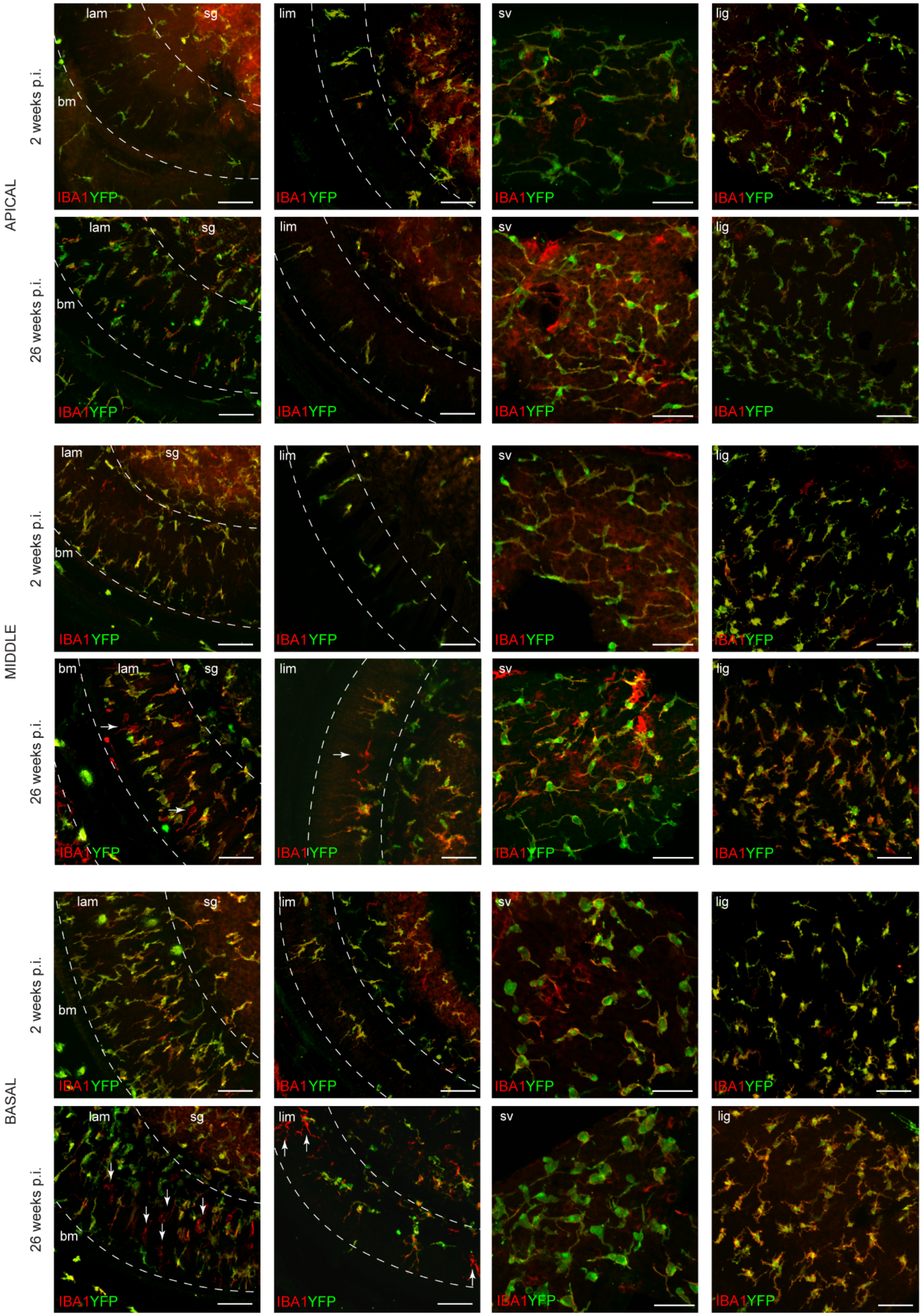
APICAL
MIDDLE
BASAL
2 weeks p.i.
26 weeks p.i.
lam
sg
bm
lim
sv
lig
IBA1YFP

Fig. S9

**Figure S9: YFP expression in IBA1+ macrophages in the apical, middle, and basal turns of the different cochlea compartments.**

Representative immunofluorescence images from the apical, middle, and basal segments of the basilar membrane (bm), spiral lamina (lam), spiral ganglion (sg), spiral limbus (lim), stria vascularis (sv), and spiral ligament (lig) from whole mounts of *$Cx3cr1^{CreERT2}$:Rosa26-YFP* mice at 2- and 26-weeks post-injection. $YFP^{-}IBA1^{+}$ single-positive cells are labeled by arrows. Representative pictures of n=11 (2 and 26 weeks each) animals. Scale bars = 50 µm.

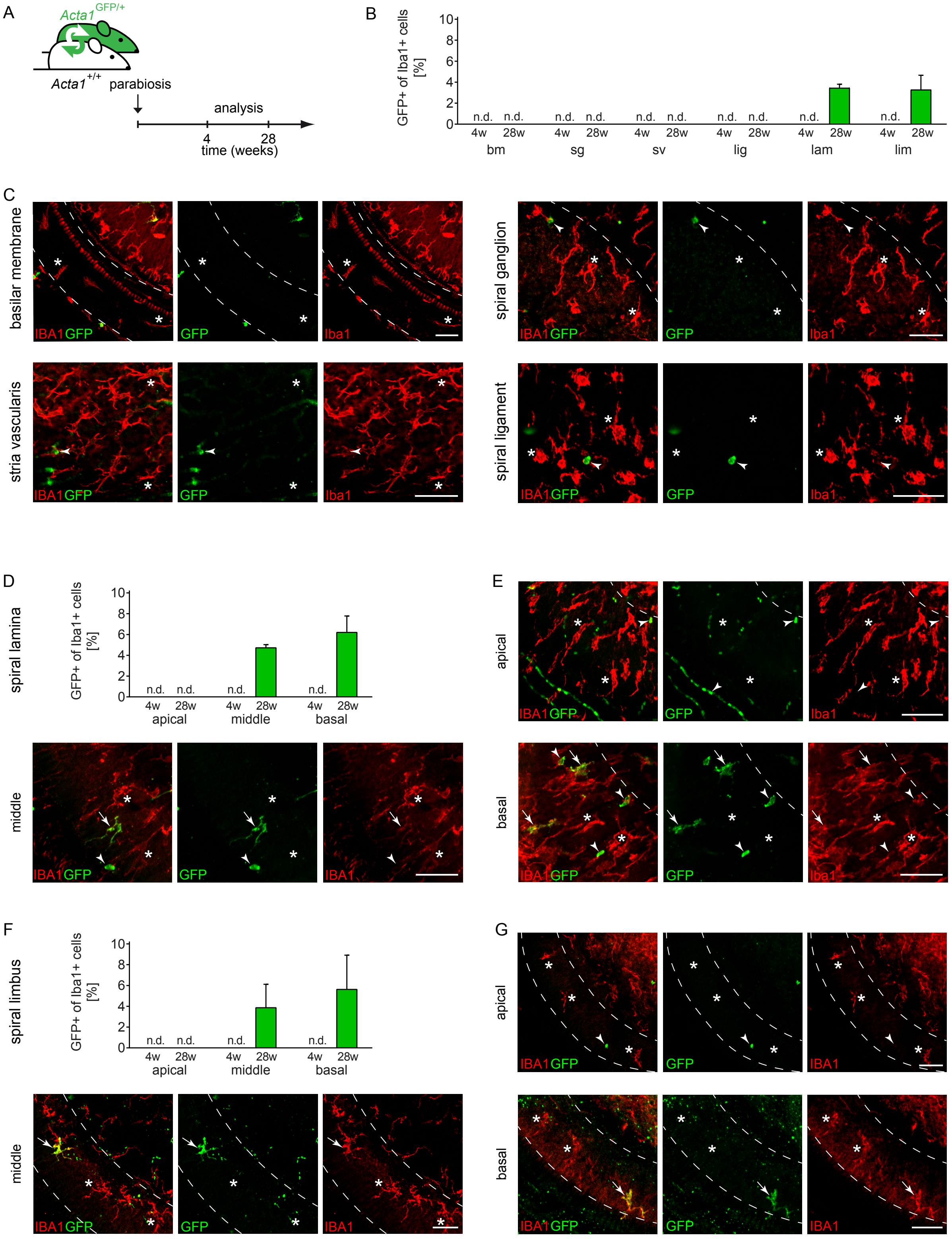
A
Acta1 GFP/+
Acta1 +/+
parabiosis
analysis
4
28
time (weeks)
B
GFP+ of Iba1+ cells [%]
n.d.
4w
28w
bm
sg
sv
lig
lam
lim
C
basilar membrane
spiral ganglion
stria vascularis
spiral ligament
IBA1GFP
GFP
Iba1
D
spiral lamina
apical
middle
basal
E
F
spiral limbus
G
IBA1

fig. S10

**Figure S10: Analysis of the contribution of blood monocytes to the cochlear macrophage pool in steady-state conditions, using parabiotic mice.**

**A**. Experimental setup of surgically connected parabiotic mice. *Ubc*-GFP and C57BL/6J mice underwent parabiosis surgery and were analysed at 4 and 28 weeks after parabiosis.

**B**. Percentage of $GFP^{+}IBA1^{+}$ macrophages among the $IBA1^{+}$ macrophages in the basilar membrane (bm), spiral lamina (lam), spiral ganglion (sg), spiral limbus (lim), stria vascularis (sv), and spiral ligament (lig). n.d. = not detectable. Bars represent mean ± s.e.m. n=4 (4 weeks) and n=4 (28 weeks).

**C**. Representative immunofluorescence images from the basilar membrane, spiral ganglion, stria vascularis, and spiral ligament from whole mounts of the non-GFP partner at 28- weeks post parabiosis. $GFP^{-}IBA1^{+}$ resident macrophages are marked (asterisks) and $GFP^{+}IBA1^{+}$ are macrophages originating from blood monocytes (arrows). Scale bars = 50 µm.

**D**. Percentage of $GFP^{+}IBA1^{+}$ macrophages among the $IBA1^{+}$ macrophages in the spiral lamina apical, middle, and basal turns. Bars represent means ± s.e.m., n=4 (4 weeks) and n=4 (28 weeks).

**E**. Representative immunofluorescence images from the apical, middle, and basal turns of the spiral lamina of the non-GFP partner at 28- weeks post parabiosis. $GFP^{-}IBA1^{+}$ resident macrophages are marked (asterisks) and $GFP^{+}IBA1^{+}$ are macrophages originating from blood monocytes (arrows). Scale bars = 50 µm.

**F**. Percent of $GFP^{+}IBA1^{+}$ macrophages among the $IBA1^{+}$ macrophages in the spiral limbus apical, middle, and basal turns. Bars represent mean ± s.e.m., n=4 (4 weeks) and n=4 (28 weeks).

**G**. Representative immunofluorescence images from the apical (l), middle (m), and basal (n) turns of the spiral limbus at 28 weeks post-surgery. $GFP^{-}IBA1^{+}$ resident macrophages are marked (asterisks) and $GFP^{+}IBA1^{+}$ are macrophages originating from blood monocytes (arrows). Scale bars = 50 µm.

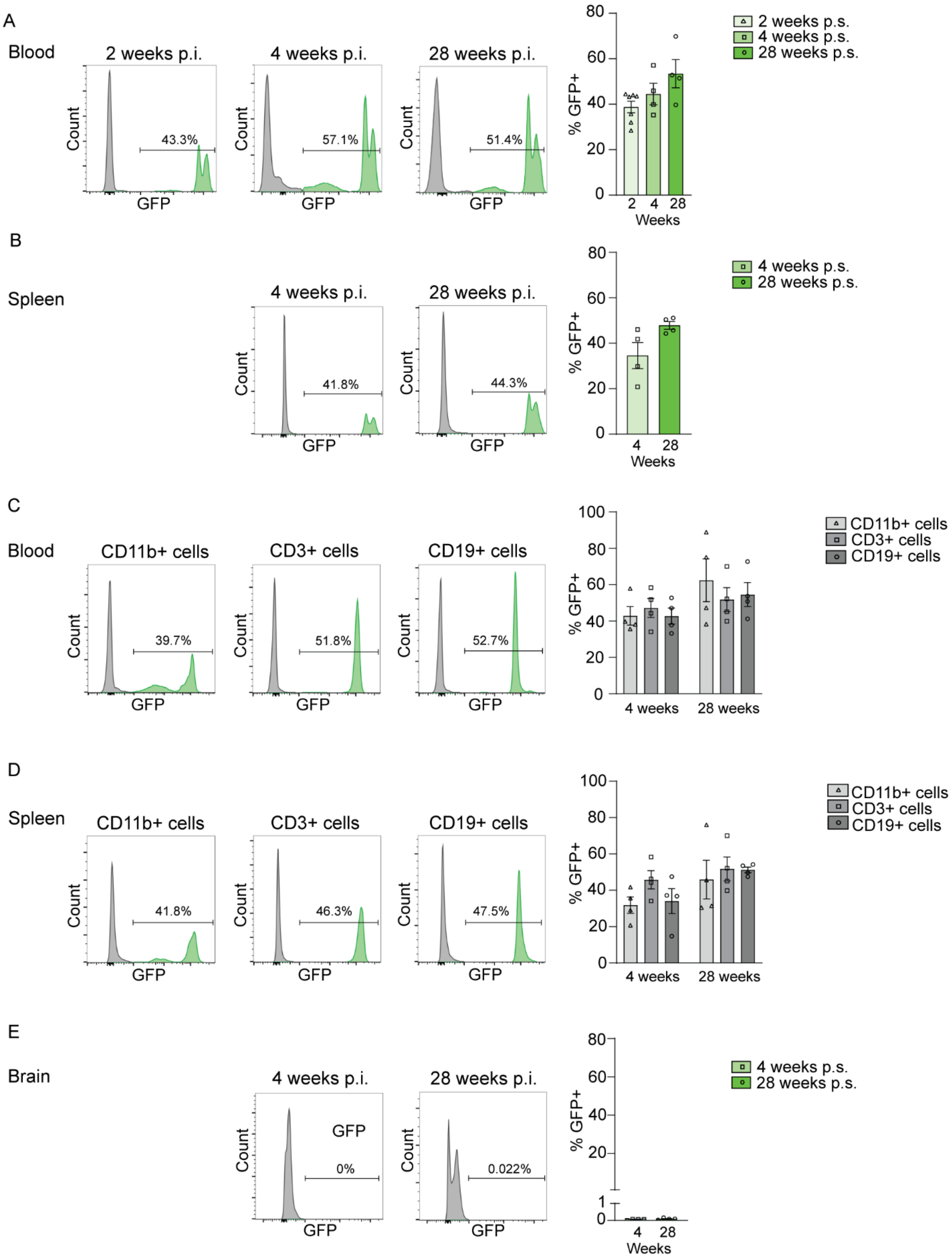
A
Blood
2 weeks p.i.
4 weeks p.i.
28 weeks p.i.
Count
GFP
43.3%
57.1%
51.4%
% GFP+
Weeks
2 weeks p.s.
4 weeks p.s.
28 weeks p.s.
B
Spleen
4 weeks p.i.
28 weeks p.i.
41.8%
44.3%
C
Blood
CD11b+ cells
CD3+ cells
CD19+ cells
39.7%
51.8%
52.7%
4 weeks
28 weeks
D
Spleen
41.8%
46.3%
47.5%
E
Brain
0%
0.022%

fig. S11

**Figure S11. GFP expression in tissues of parabiosis mice.**

**A-B.** Histogram showing the expression of GFP by $CD11b^+$ blood myeloid cells (a) and $CD11b^+$ splenocytes (b) in the non-GFP wildtype partner of the parabiosis pair at 2, 4, and 28 weeks post parabiosis surgery (p.s.). Quantification of the percentage of $CD11b^+$ blood cells expressing GFP in the non-GFP partner 2, 4, and 28 weeks p.s.

**C-D.** Histogram showing a representative expression of GFP in $CD11b^+$, $CD3^+$ T cells, and $CD19^+$ B cells from the blood (c) and the spleen (d) of the non-GFP partner at 28 weeks post parabiosis. Quantification of the percentage of blood GFP-expressing $CD11b^+$, $CD3^+$ T cells, and $CD19^+$ B cells 4 and 28 weeks p.s.

**E.** Histogram showing the expression of GFP by brain microglial cells of the non-GFP partner of the parabiosis pair 4, and 28 weeks p.s. Quantification of the percentage of $CD11b^+$ myeloid cells expressing GFP in the non-GFP partner pair 4 and 28 weeks p.s.

Data are from three independent experiments with n=7 (2 weeks p.s.), n=4 mice (4 weeks), and n=4 mice (28 weeks) and presented as mean ± s.e.m.

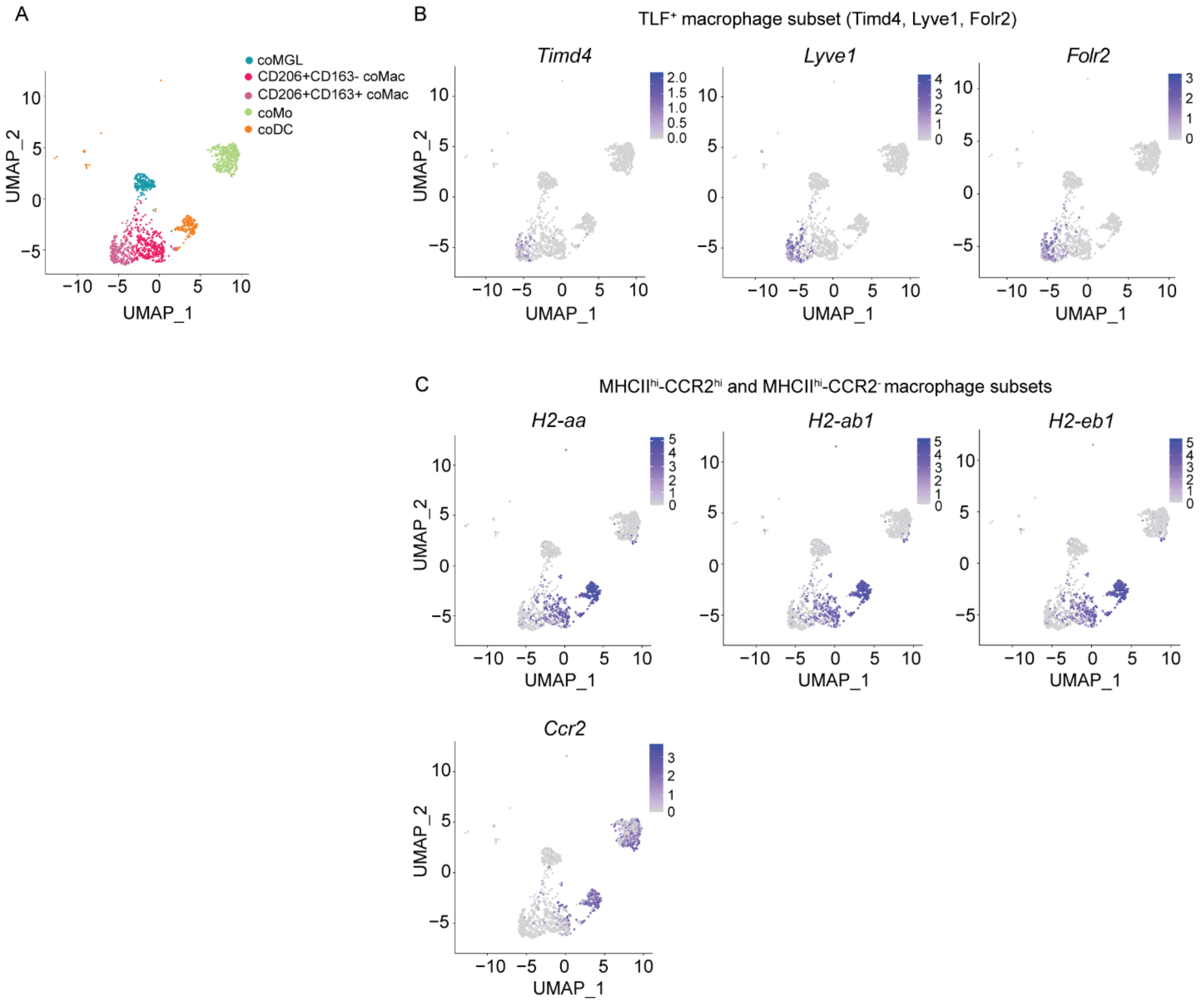


fig. S12

**Figure S12. TLF$^{+}$/Lyve1$^{hi}$, MHCII$^{hi}$-Ccr2$^{hi}$ and MhcII$^{hi}$-Ccr2$^{-}$ macrophage subset markers expression in the cochlea.**

**A.** UMAP plot (from Figure1a) depicting five distinct clusters (cluster α to ε).

**B.** UMAP plot depicting the expression of *Timd4*, *Lyve1* and *Folr2*, markers of the TLF$^{+}$/Lyve1$^{hi}$ macrophage subset previously reported in the literature (*51,52*).

**C.** UMAP plot depicting the expression of MHC-II related genes (*H2-aa*, *H2-ab1*, *H2-eb1*) and *Ccr2*, markers of the MHCII$^{hi}$-CCR2$^{hi}$ and MHCII$^{hi}$-CCR2$^{-}$ macrophage subsets previously reported in the literature (*51,52*).

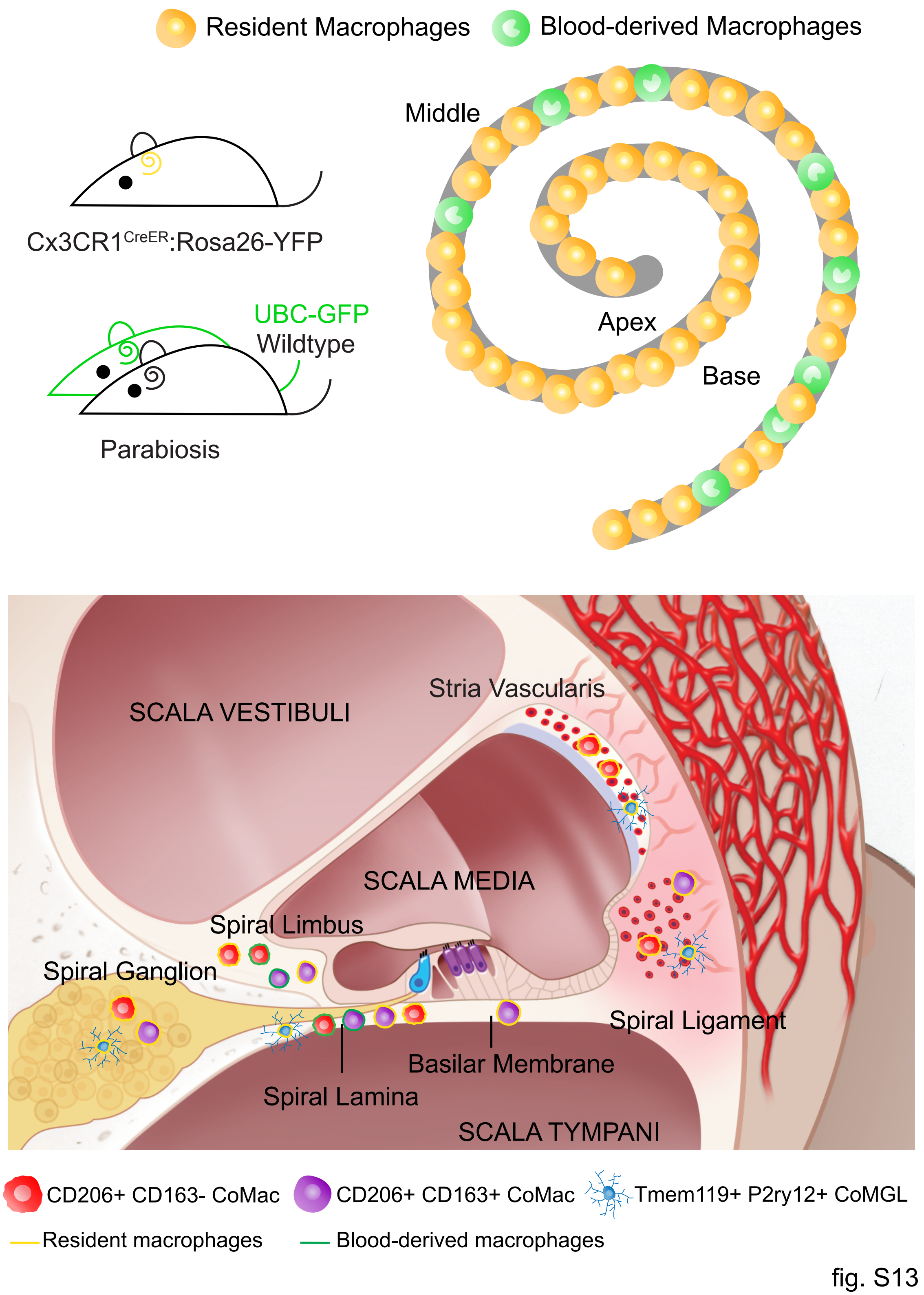


166

167 **Figure S13. Graphical abstract of experiment findings.**

| Antibodies | | |
| --- | --- | --- |
| **Antibody, dilution** | **Manufacturer** | **Catalog #** |
| rabbit anti-Iba-1, polyclonal, 1:500 | Synaptic Systems, Göttingen, Germany | 234 013 |
| rabbit anti mouse-Iba-1, 1:500 | Fujifilm Wako Chemicals, USA | 019-19741 |
| guinea pig anti-Iba1 (clone Gp311H9), monoclonal, 1:500 | Synaptic Systems, Göttingen, Germany | HS-234 308 |
| goat anti-GFP, polyclonal, 1:200 | OriGene Technologies, Rockville, USA | R1091P |
| chicken anti-GFP, 1:800 | Thermo Fisher Scientific Inc., USA | A10262 |
| rabbit anti-P2RY12, polyclonal, 1:100 | AnaSpec, Fremont, USA | AS-55043A |
| rabbit anti-Tmem119, polyclonal, 1:500 | Synaptic Systems, Göttingen, Germany | 400 002 |
| rat anti-CD206-Alexa Fluor 647 (clone MR5D3), monoclonal, 1:100 | Bio-Rad Laboratories, UK | MCA2235A647 |
| rat anti mouse-CD206, 1:100 | Bio-Rad Laboratories, UK | MCA2235GA |
| rat anti-CD163 (clone S15049I), monoclonal, 1:100 | BioLegend, USA | 155302 |
| goat anti-rabbit, Alexa Fluor 488, polyclonal, 1:500 | Thermo Fisher Scientific Inc., USA | A11034 |
| donkey anti-rabbit, Alexa Fluor 555, 1:500 | Thermo Fisher Scientific Inc., USA | A31572 |
| donkey anti-rabbit, Alexa Fluor 568, polyclonal, 1:500 | Thermo Fisher Scientific Inc., USA | A10042 |
| goat anti-rabbit, Alexa Fluor 647, polyclonal, 1:500 | Thermo Fisher Scientific Inc., USA | A21245 |
| donkey anti-rabbit, Alexa Fluor 647, 1:500 | Thermo Fisher Scientific Inc., USA | A31573 |
| goat anti-guinea pig, Alexa Fluor 568, polyclonal, 1:500 | Thermo Fisher Scientific Inc., USA | A11075 |
| goat anti-guinea pig, DyLight 594, 1:500 | Thermo Fisher Scientific Inc., USA | SA5-10096 |
| donkey anti-goat, Alexa Fluor 647, polyclonal, 1:500 | Thermo Fisher Scientific Inc., USA | A21447 |
| goat anti-chicken, Alexa Fluor 488, 1:500 | Thermo Fisher Scientific Inc., USA | A11039 |
| goat anti-rat, Alexa Fluor 568, polyclonal, 1:500 | Thermo Fisher Scientific Inc., USA | A11077 |
| donkey anti-rat, Alexa Fluor 594, 1:500 | Thermo Fisher Scientific Inc., USA | A21209 |

168 **table S1. List of antibodies.**

| | TAM (n=7) | | untreated (n=3) | | |
|---|---|---|---|---|---|
| **compartment** | **mean** | **SEM** | **mean** | **SEM** | **p-value** |
| **basilar membrane** | 8.45 | 0.63 | 1.80 | 0.16 | <0.0001 |
| **spiral lamina** | 5.85 | 0.51 | 1.89 | 0.49 | 0.0035 |
| **spiral ganglion** | 8.40 | 1.11 | 1.68 | 0.48 | 0.0011 |
| **spiral limbus** | 3.98 | 0.76 | 1.59 | 0.65 | 0.0813 |
| **stria vascularis** | 9.94 | 0.63 | 2.52 | 0.75 | 0.0021 |
| **spiral ligament** | 11.92 | 1.12 | 4.57 | 0.67 | 0.0011 |

**table S2. Summary table of values for pulse labeling experiment (Figure S8)**

| | 2 wpi (n=11) | | 8 wpi (n=6) | | 26 wpi (n=11) | | |
|---|---|---|---|---|---|---|---|
| **compartment** | **mean** | **SEM** | **mean** | **SEM** | **mean** | **SEM** | **p-value** |
| **basilar membrane** | 97.03 | 0.77 | 96.39 | 1.21 | 93.79 | 1.37 | 0.1995 |
| **spiral lamina** | 94.53 | 2.10 | 77.64 | 2.59 | 78.25 | 1.50 | 0.0001 |
| **spiral ganglion** | 95.01 | 1.58 | 92.60 | 4.37 | 91.40 | 2.23 | 0.4832 |
| **spiral limbus** | 93.69 | 1.49 | 81.81 | 3.78 | 77.63 | 3.99 | 0.0059 |
| **stria vascularis** | 95.59 | 1.26 | 94.85 | 2.92 | 94.09 | 1.77 | 0.9880 |
| **spiral ligament** | 99.63 | 0.18 | 100.66 | 2.19 | 97.85 | 0.77 | 0.1778 |

| **basilar membrane** | 2 wpi (n=11) | | 8 wpi (n=6) | | 26 wpi (n=11) | | |
|---|---|---|---|---|---|---|---|
| **level** | **mean** | **SEM** | **mean** | **SEM** | **mean** | **SEM** | **p-value** |
| **apical** | 96.52 | 0.92 | 93.55 | 3.43 | 95.40 | 1.47 | 0.6773 |
| **middle** | 96.50 | 1.39 | 97.32 | 1.24 | 94.28 | 2.06 | 0.8546 |
| **basal** | 96.93 | 0.87 | 95.90 | 1.59 | 91.12 | 3.96 | 0.8011 |
| **p-value** | 0.5742 | | 0.5261 | | 0.5250 | | |

| **spiral lamina** | 2 wpi (n=11) | | 8 wpi (n=6) | | 26 wpi (n=11) | | |
|---|---|---|---|---|---|---|---|
| **level** | **mean** | **SEM** | **mean** | **SEM** | **mean** | **SEM** | **p-value** |
| **apical** | 96.88 | 1.62 | 88.12 | 3.11 | 91.07 | 2.16 | 0.0599 |
| **middle** | 95.11 | 2.02 | 78.17 | 3.74 | 79.59 | 1.86 | 0.0004 |
| **basal** | 91.85 | 3.30 | 69.69 | 3.42 | 66.41 | 1.67 | 0.0001 |
| **p-value** | 0.0568 | | 0.0119 | | <0.0001 | | |

| **spiral ganglion** | 2 wpi (n=11) | | 8 wpi (n=6) | | 26 wpi (n=11) | | |
|---|---|---|---|---|---|---|---|
| **level** | **mean** | **SEM** | **mean** | **SEM** | **mean** | **SEM** | **p-value** |
| **apical** | 94.20 | 2.33 | 93.03 | 2.80 | 94.89 | 1.54 | 0.8696 |
| **middle** | 95.94 | 2.07 | 95.86 | 5.79 | 92.05 | 2.57 | 0.5546 |
| **basal** | 93.01 | 1.30 | 90.68 | 4.74 | 87.80 | 3.32 | 0.8381 |
| **p-value** | 0.4204 | | 0.4372 | | 0.2557 | | |

| **spiral limbus** | 2 wpi (n=11) | | 8 wpi (n=6) | | 26 wpi (n=11) | | |
|---|---|---|---|---|---|---|---|
| **level** | **mean** | **SEM** | **mean** | **SEM** | **mean** | **SEM** | **p-value** |
| **apical** | 94.10 | 2.53 | 89.54 | 4.93 | 91.13 | 3.18 | 0.7258 |
| **middle** | 94.74 | 1.59 | 83.27 | 6.51 | 79.36 | 4.11 | 0.0106 |
| **basal** | 92.43 | 2.15 | 78.16 | 3.91 | 69.35 | 4.08 | 0.0010 |
| **p-value** | 0.7171 | | 0.2920 | | <0.0001 | | |

| **stria vascularis** | 2 wpi (n=11) | | 8 wpi (n=6) | | 26 wpi (n=11) | | |
|---|---|---|---|---|---|---|---|
| **level** | **mean** | **SEM** | **mean** | **SEM** | **mean** | **SEM** | **p-value** |
| **apical** | 93.67 | 1.41 | 96.84 | 5.19 | 92.29 | 2.59 | 0.7046 |
| **middle** | 96.81 | 2.20 | 93.80 | 2.19 | 93.32 | 1.68 | 0.5000 |
| **basal** | 96.30 | 1.26 | 96.75 | 3.92 | 94.81 | 2.35 | 0.9731 |
| **p-value** | 0.3088 | | 0.7330 | | 0.4694 | | |

| **spiral ligament** | 2 wpi (n=11) | | 8 wpi (n=6) | | 26 wpi (n=11) | | |
|---|---|---|---|---|---|---|---|
| **level** | **mean** | **SEM** | **mean** | **SEM** | **mean** | **SEM** | **p-value** |
| **apical** | 99.72 | 0.90 | 101.38 | 2.93 | 98.56 | 0.63 | 0.6486 |
| **middle** | 99.89 | 0.86 | 100.94 | 2.11 | 98.50 | 0.91 | 0.2054 |
| **basal** | 98.94 | 0.24 | 100.20 | 2.32 | 97.02 | 1.17 | 0.4934 |
| **p-value** | 0.0685 | | >0.9999 | | 0.4026 | | |

**table S3. Summary table of values for turn-over experiment (Figure 4)**

| | 4 wps (n=4) | | 28 wps (n=4) | | |
|---|---|---|---|---|---|
| compartment | mean | SEM | mean | SEM | p-value |
| basilar membrane | 0.00 | 0.00 | 0.00 | 0.00 | n.a. |
| spiral lamina | 0.00 | 0.00 | 3.46 | 0.30 | n.a. |
| spiral ganglion | 0.00 | 0.00 | 0.00 | 0.00 | n.a. |
| spiral limbus | 0.00 | 0.00 | 3.29 | 1.19 | n.a. |
| stria vascularis | 0.00 | 0.00 | 0.00 | 0.00 | n.a. |
| spiral ligament | 0.00 | 0.00 | 0.00 | 0.00 | n.a. |

| basilar membrane | 4 wps (n=4) | | 28 wps (n=4) | | |
|---|---|---|---|---|---|
| level | mean | SEM | mean | SEM | p-value |
| apical | 0.00 | 0.00 | 0.00 | 0.00 | n.a. |
| middle | 0.00 | 0.00 | 0.00 | 0.00 | n.a. |
| basal | 0.00 | 0.00 | 0.00 | 0.00 | n.a. |
| p-value | n.a. | | n.a. | | |

| spiral lamina | 4 wps (n=4) | | 28 wps (n=4) | | |
|---|---|---|---|---|---|
| level | mean | SEM | mean | SEM | p-value |
| apical | 0.00 | 0.00 | 0.00 | 0.00 | n.a. |
| middle | 0.00 | 0.00 | 4.72 | 0.27 | n.a. |
| basal | 0.00 | 0.00 | 6.20 | 1.37 | n.a. |
| p-value | n.a. | | n.a. | | |

| spiral ganglion | 4 wps (n=4) | | 28 wps (n=4) | | |
|---|---|---|---|---|---|
| level | mean | SEM | mean | SEM | p-value |
| apical | 0.00 | 0.00 | 0.00 | 0.00 | n.a. |
| middle | 0.00 | 0.00 | 0.00 | 0.00 | n.a. |
| basal | 0.00 | 0.00 | 0.00 | 0.00 | n.a. |
| p-value | n.a. | | n.a. | | |

| spiral limbus | 4 wps (n=4) | | 28 wps (n=4) | | |
|---|---|---|---|---|---|
| level | mean | SEM | mean | SEM | p-value |
| apical | 0.00 | 0.00 | 0.00 | 0.00 | n.a. |
| middle | 0.00 | 0.00 | 3.87 | 1.95 | n.a. |
| basal | 0.00 | 0.00 | 5.63 | 2.85 | n.a. |
| p-value | n.a. | | n.a. | | |

| stria vascularis | 4 wps (n=4) | | 28 wps (n=4) | | |
|---|---|---|---|---|---|
| level | mean | SEM | mean | SEM | p-value |
| apical | 0.00 | 0.00 | 0.00 | 0.00 | n.a. |
| middle | 0.00 | 0.00 | 0.00 | 0.00 | n.a. |
| basal | 0.00 | 0.00 | 0.00 | 0.00 | n.a. |
| p-value | n.a. | | n.a. | | |

| spiral ligament | 4 wps (n=4) | | 28 wps (n=4) | | |
|---|---|---|---|---|---|
| level | mean | SEM | mean | SEM | p-value |
| apical | 0.00 | 0.00 | 0.00 | 0.00 | n.a. |
| middle | 0.00 | 0.00 | 0.00 | 0.00 | n.a. |
| basal | 0.00 | 0.00 | 0.00 | 0.00 | n.a. |
| p-value | n.a. | | n.a. | | |

**table S4. Summary table of values for parabiosis experiment (Figure S10)**

**Iba1+ cells per 200um x 200um**

| | 7w (n=3) | | 36w (n=3) | | 70w (n=3) | | p-value | | |
|---|---|---|---|---|---|---|---|---|---|
| compartment | mean | SEM | mean | SEM | mean | SEM | 7w-36w | 36-70w | 7w-70w |
| spiral lamina | 10.00 | 0.24 | 11.83 | 0.36 | 15.50 | 0.24 | 0.0322 | 0.0038 | 0.0002 |
| spiral ganglion | 8.67 | 0.27 | 9.33 | 0.27 | 9.00 | 0.47 | 0.6000 | >0.9999 | >0.9999 |
| stria vascularis | 7.00 | 0.82 | 6.17 | 0.14 | 7.50 | 1.18 | >0.9999 | 0.6000 | >0.9999 |
| spiral ligament | 12.33 | 0.95 | 18.00 | 1.41 | 20.33 | 0.54 | 0.0616 | 0.4000 | 0.1000 |

**CD206+ of Iba1+ cells**

| | 7w (n=3) | | 36w (n=3) | | 70w (n=3) | | p-value | | |
|---|---|---|---|---|---|---|---|---|---|
| compartment | mean | SEM | mean | SEM | mean | SEM | 7w-36w | 36-70w | 7w-70w |
| spiral lamina | 51.41 | 4.51 | 60.23 | 6.41 | 83.55 | 1.68 | 0.4259 | 0.1027 | 0.0320 |
| spiral ganglion | 55.15 | 6.50 | 65.07 | 1.74 | 73.97 | 4.70 | 0.3382 | 0.2581 | 0.1346 |
| stria vascularis | 41.29 | 12.17 | 48.41 | 10.06 | 34.10 | 1.39 | 0.7321 | 0.3651 | 0.6776 |
| spiral ligament | 43.54 | 1.90 | 47.49 | 10.20 | 54.50 | 3.25 | 0.7834 | 0.6381 | 0.0915 |

**TMEM119+ of Iba1+ cells**

| | 7w (n=3) | | 36w (n=3) | | |
|---|---|---|---|---|---|
| compartment | mean | SEM | mean | SEM | p-value |
| spiral lamina | 36.21 | 4.19 | 40.42 | 1.99 | 0.5150 |
| spiral ganglion | 14.24 | 0.12 | 15.30 | 2.50 | 0.7612 |
| stria vascularis | 19.09 | 1.30 | 21.91 | 4.33 | 0.6548 |
| spiral ligament | 57.13 | 2.07 | 56.90 | 4.27 | 0.7000 |

**table S5. Summary table of values for ageing experiment (Figure 5)**
